# The Christchurch point mutation in mouse APOE reduces Aβ-induced tau and α-synuclein pathologies

**DOI:** 10.1101/2025.11.05.686857

**Authors:** Carlos M. Soto-Faguás, Abigail O’Niel, Paul A. Mueller, Paula Sanchez-Molina, Randy L. Woltjer, Jacob Raber, Vivek K. Unni

**Affiliations:** Department of Neurology Oregon Health & Science University, Portland, OR, USA; Jungers Center for Neurosciences Research, Oregon Health & Science University, Portland, OR, USA; Department of Behavioral Neuroscience, Oregon Health & Science University, Portland, OR, USA; Center for Preventive Cardiology, Knight Cardiovascular Institute, Oregon Health & Science University, Portland, OR, USA; Independent Researcher, Portland, OR, USA; Department of Pathology and Laboratory Medicine, Oregon Health & Science University, Portland, OR, USA; Department of Radiation Medicine and Division of Neuroscience, ONPRC, Oregon Health & Science University, Portland, OR, USA; OHSU Parkinson Center, Oregon Health & Science University, Portland, OR, USA

## Abstract

*Apolipoprotein E* (*APOE*) genotype is well known to influence both amyloid-β (Aβ) and tau pathologies and risk for Alzheimer’s disease (AD), but it also affects α-synuclein (α-syn) levels, Lewy pathology and risk of dementia in Parkinson’s disease (PD) and dementia with Lewy bodies (DLB). The APOE-R136S (Christchurch, CC) point mutation has been shown to protect against AD pathology and dementia, however, the molecular mechanisms underlying this protection and its effects on α-syn pathology are not well understood. Using CRISPR/Cas9 technology, we created a CC arginine-to-serine point mutation at the conserved location in mouse APOE (R128S) to understand its effects on Aβ, tau and α-syn pathologies. We crossed these APOE CC mice to 5xFAD, PS19 and A53T-αSyn-GFP (A53T) mice. Using these various double mutant mice, we tested the effect of mouse APOE CC on different proteinopathies, including Aβ, tau, Aβ-induced tau after paired helical filament (PHF)-tau intracortical injections, and α-syn after preformed fibril (PFF) intracortical and intramuscular injections. We used immunohistochemical, biochemical and behavioral measures to test for protective effects of APOE CC on these different proteinopathies. Heterozygous (Het) and homozygous (Hom) APOE CC mice showed increased plasma cholesterol and triglyceride levels, as seen in humans, but no differences in body or brain weight, or life expectancy. APOE CC decreased Aβ-induced tau pathologies in PHF-tau injected 5xFAD;Hom mice but did not change Aβ-plaque pathology in 5xFAD mice or tau pathology in PS19 mice. Although Aβ levels, tau levels and mouse sex correlated strongly with the behavioral performance, we only detected subtle effects of APOE CC on anxiety-like behaviors in crosses with 5xFAD, PS19 and PHF-tau injected 5xFAD mice. Interestingly, Het and Hom APOE CC mice both showed reduced formation and spread of Lewy pathology in brain after intracortical α-syn PFF injection and reduced formation in spinal cord after α-syn PFF injection into the hindlimb gastrocnemius muscle in A53T mice. Our study emphasizes the protective effects of the APOE CC variant against different proteinopathies important for dementia and movement disorders, including Aβ plaque, tau and α-syn, and suggests that targeting APOE CC could provide new therapeutic strategies for AD, DLB and PD.

**Graphical Abstract:** 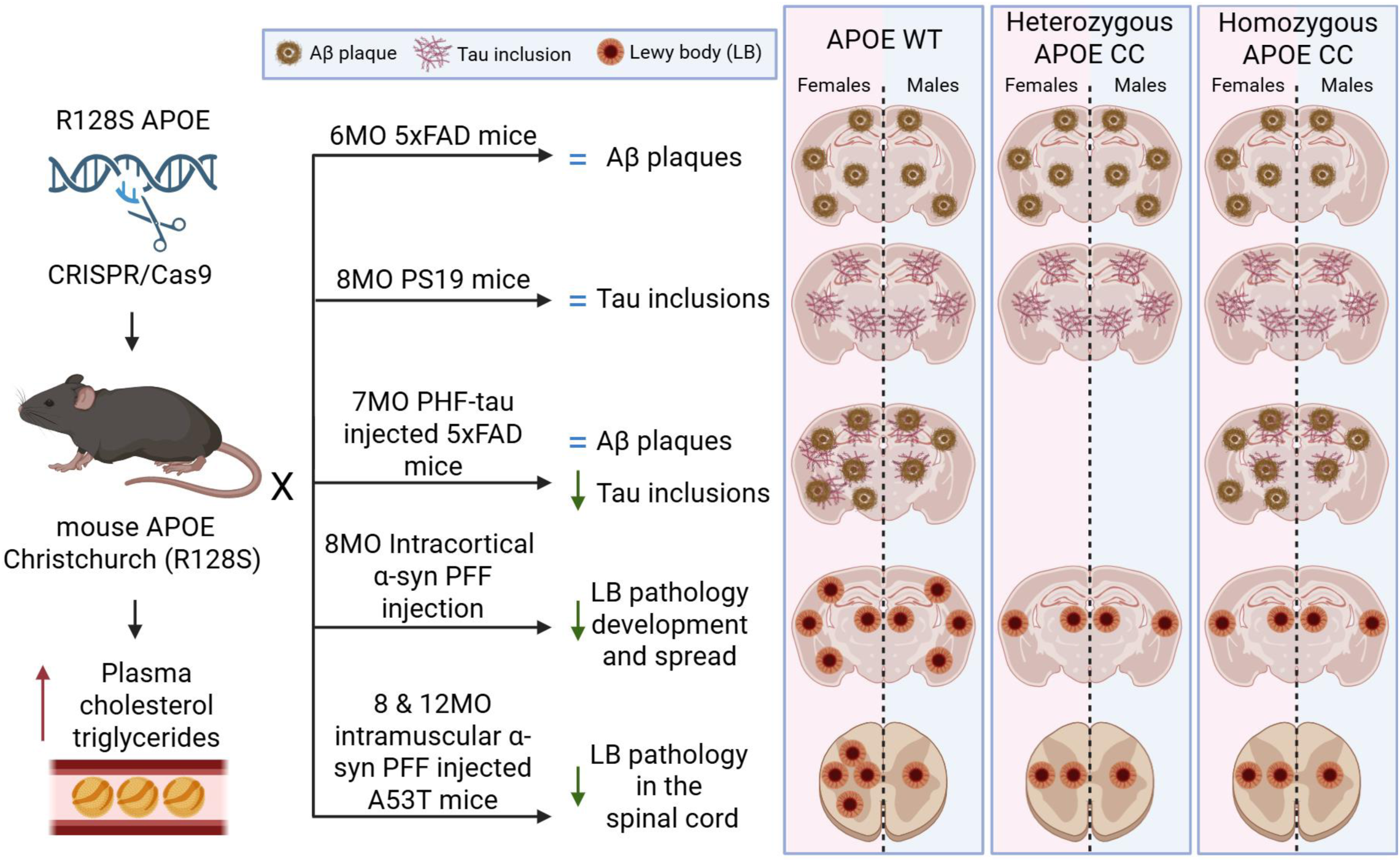

## Introduction

Lewy bodies (LB) are intracytoplasmic inclusions principally composed of aggregated α-synuclein (α-syn). They are the pathological hallmark of several clinically important neurodegenerative disorders, including Parkinson’s disease (PD), dementia with LB (DLB) and LB variant Alzheimer’s disease (LBvAD). Importantly, DLB and LBvAD patients have more rapid cognitive decline and faster progression to death than those with AD but without LB,^1,2^ suggesting a synergistic toxicity between α-syn and amyloid-β (Aβ)/tau pathology. Cholinesterase inhibitors and dopaminergic therapies can provide symptomatic benefits,^3^ however, there are no currently approved therapies that slow or halt progressive LB-induced neurodegeneration.

Genetic studies have highlighted the importance of *Apolipoprotein E (APOE)* genotype in multiple neurodegenerative disorders.^4,5,6^ APOE is the most abundant apolipoprotein in the brain, and under physiological conditions mainly produced by astrocytes, but in response to stress can also be produced by neurons, microglia and oligodendrocytes.^7,8,9,10^ Mature human APOE is secreted as a 299 amino acid protein with three different domains: an N-terminal receptor-binding domain, a central linker, and a C-terminal region implicated in lipid binding.^11^ There are three common alleles in humans: *APOE2*, *APOE3* and *APOE4* which encode for APOE2, APOE3 and APOE4 isoforms, respectively. These variants show different protein structure and function, despite differing by only two amino acids at positions 112 and 158, with APOE2 having two cysteines, APOE3 one cysteine and one arginine, and APOE4 two arginines, respectively. Conversely, mice have only one APOE isoform that has two arginines at the conserved positions 104 and 150, more resembling the human APOE4 protein. However, mouse APOE has a threonine at position 53, instead of an arginine at the conserved position 61 found in humans, suggesting that in some ways it could also behave more like human APOE3.^12,13^

In AD, the *APOE2* allele provides protection relative to the more common *APOE3* allele, while *APOE4* confers susceptibility for developing AD.^4^ Although less well studied, recent reports also suggest a correlation between *APOE* genotype with PD pathology progression and dementia risk in PD and DLB.^5^ Consistent with this, *APOE4* correlates with higher phosphorylated and aggregated α-syn levels.^6^ Interestingly, a recent report found an individual with the autosomal dominant familial AD mutation PSEN1-E280A who was resistant to dementia.^14^ Brain PET analysis showed high Aβ but lower tau levels in specific brain areas compared to other PSEN1-E280A mutation carriers, and this was confirmed in subsequent postmortem studies.^15^ Whole genome sequencing in this woman revealed a rare homozygous R136S (“Christchurch”, CC) mutation in the *APOE3* gene, previously known to cause type III hyperlipidemia.^14,16^ This PSEN1-E280A/APOE CC patient exhibited reduced neuronal loss, glial and vascular abnormalities than expected, despite having high Aβ burden.^17^ A distinct pathological profile showing severe tau neuropathology, cerebral amyloid angiopathy and chronic inflammatory microglial responses was observed in the occipital cortex, a region with lower *APOE* expression, whereas the frontal cortex, a brain region with higher *APOE* expression, was relatively spared from tau pathology and glial alterations.^15^ These findings suggest a protective role of APOE CC against the formation of tau pathology and dementia, even in the presence of a normally fully penetrant familial AD mutation.

Given excitement around the possible protective effects of APOE CC in AD-related neurodegeneration, several models have been generated to test this further.^18,19,20,21,22,23,24,25,26^ Several of these reports show that APOE CC also reduces AD-related Aβ and tau pathologies in mice. Human *APOE* with the CC mutation knock in (KI) has been shown to reduce Aβ pathology and tau pathology in Aβ-forming^19,20,24^ and tauopathy mouse models.^18,19,20,24,25^ *In vitro* studies also showed that APOE CC’ had a protective effect against tau pathology.^21,22,23^ KI of the CC mutation into mouse *Apoe* also showed reduced Aβ-plaque, but not changes in tau pathology, when crossed with Aβ- and tau-forming mice, respectively.^26^ Interestingly, several of these studies suggest an important role for microglia^18,19,,23,25^ and astrocytes^18,21^ in the protective effects mediated by *APOE CC*. Although the mechanisms involved are not completely understood, several have been suggested. These include weaker binding to heparan sulfate proteoglycans (HSPG),^14,18,19,27^ direct binding to tau reducing its cellular uptake, fragmentation and spread,^24^ alterations in microglial and astrocytic responses,^18,19,20,21^ upregulation of cadherin and WNT/β-catenin signaling,^22,28^ reduction of microglial cGAS-STING-IFN pathway activation,^25^ and changes in microglial lipid peroxidation and phagocytosis.^23^ In contrast to what has been reported for AD-related Aβ and tau pathology, to our knowledge, the possible role of APOE CC in modulating α-syn LB pathology has not been studied to date.

Given the very low frequency of the APOE CC mutation, it is difficult to study its potential protective effects in different forms of neurodegeneration in humans. Thus, a mouse model recapitulating these phenotypes would be useful to understand whether and how APOE CC protects against different proteinopathies. Although the human and mouse APOE protein share ∼70% homology overall, the receptor-binding domain is even more highly conserved (∼85% homology) and is implicated in APOE binding to HSPG and LDLR, which may directly modulate the formation of AD-like pathology.^14.19^ Interestingly, the R136 residue in human APOE that is mutated in CC is conserved in mice, corresponding to residue R128. Given this interest in understanding the role of APOE CC in multiple proteinopathies, we have developed a mouse model expressing the murine APOE CC (R128S) mutation and crossed it with the Aβ-plaque forming 5xFAD mice, the tau tangle-forming PS19 mice, and the A53T α-syn-overexpressing α Syn-GFP (A53T) mice to study the role of this murine variant in Aβ, tau and α-syn pathology formation and spread. Here, we describe the role of murine APOE CC in the development of Aβ-induced tau tangles and, for the first time, LB pathology in the brain and in LB peripheral to central nervous system (CNS) spread.

## Results

### Generation and characterization of murine APOE CC mice

Given that human *APOE* KI into the mouse locus delays Aβ pathology formation on its own^29^ and to avoid potential confounds associated with altered interactions between human APOE with different murine receptors, we decided to generate the murine APOE R128S CC allele by using CRISPR/Cas9 technology (**Figure 1A**).

**Figure 1.**
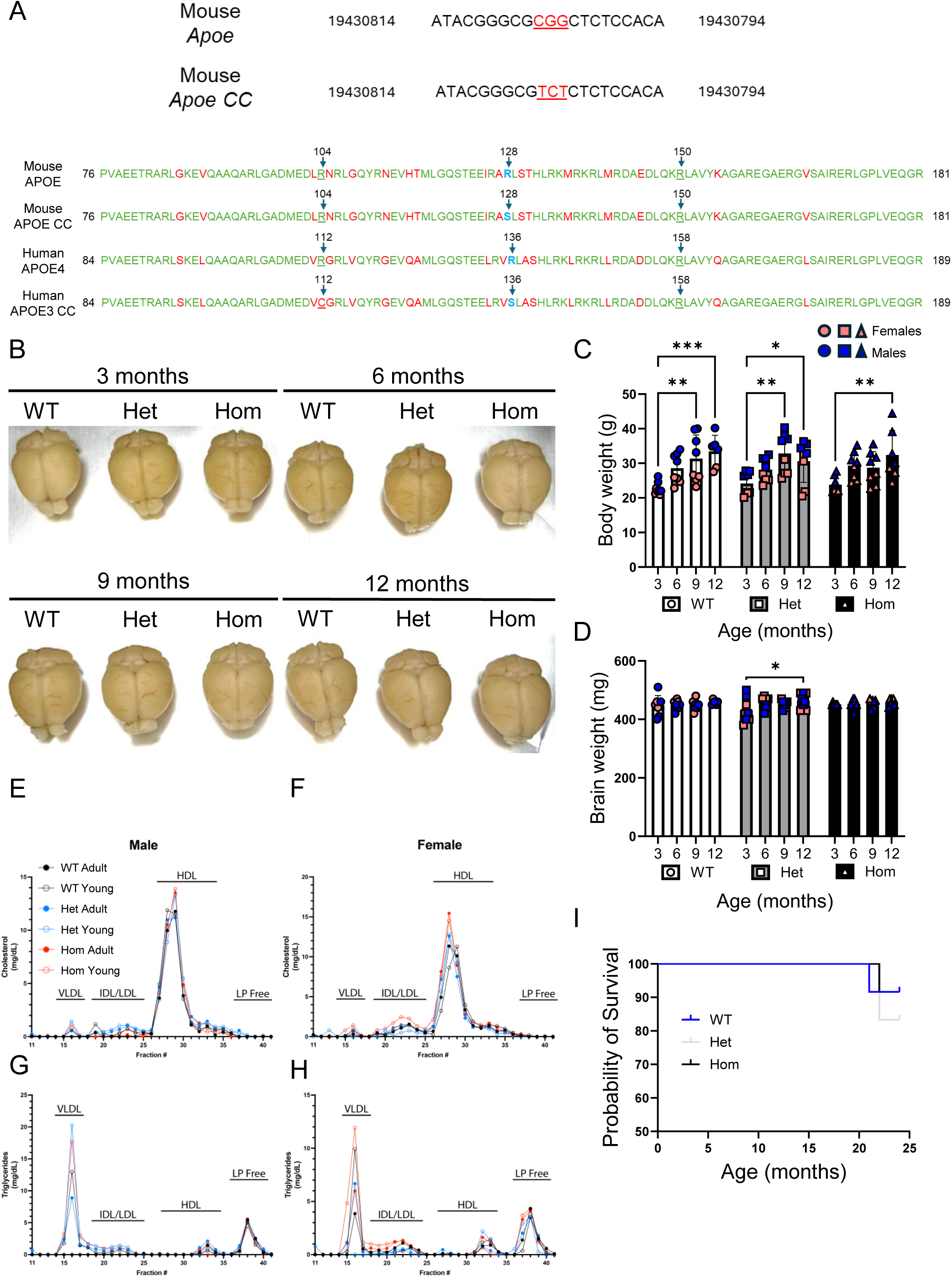
Generation and characterization of the murine APOE CC mouse. **A** Targeted codon in the mouse APOE gene to generate the murine APOE CC mutation. Partial protein sequence comparison between mouse APOE, mouse APOE CC, human APOE4 and human APOE3 CC. **B** Representative whole brain images from 3, 6, 9 and 12 months-old WT, Het and Hom mice. **C-D** Quantification of the body [**C**, age effect: F (3,80) = 14.22, *p* < 0.0001] and brain (**D**) weights. Data represent mean ± standard deviation (SD) of multiple mice (n = 4/genotype, sex and age). Two-way ANOVA followed by Tukey’s post hoc test was used as statistical test. * *p* < 0.05, ** *p* < 0.01, *** *p* < 0.001. **E-H** FPLC analysis of the plasma lipoproteins in young (<2 months) and adult (4 months) mice. 150 µl of pooled plasma from three mice from the same group (50 µl each) was used. **I** Life expectancy analysis of mice up to 24 months of age (n = 12 mice/group). Mantel-Cox test was used as statistical test.

To investigate the effects of murine APOE CC in the heterozygous (Het) or homozygous (Hom) state compared to control littermates (WT), we measured the body and brain weights of 3, 6, 9 and 12 month-old mice (**Figure 1B-D**). Body weight measurements revealed an expected age effect with increased weight in 9-and 12-month-old WT and Het and 12-month-old Hom mice compared to their respective 3-month-old group, without significant differences between genotypes (**Figure 1C**). Similarly, no differences were found between genotypes in the brain weight across these ages (**Figure 1D**). We detected an increase in brain weight in 12-month-old Het when compared to 3-month-old Het mice (**Figure 1D**).

The APOE CC mutation has been previously shown to interfere with receptor-mediated-uptake of APOE containing lipoprotein particles, leading to type III hyperlipoproteinemia.^16^ We used fast protein liquid chromatography (FPLC) to analyze plasma cholesterol and triglyceride levels in young (<2 months) and adult (4 months) WT, Het and Hom APOE CC mice. Lipid analysis showed increased very low-density lipoprotein (VLDL)- and high-density lipoprotein (HDL)-cholesterol in young Het and Hom males and adult Hom males compared to WT males (**Figure 1E**). Similarly, young and adult Het and Hom females showed increased HDL-cholesterol levels compared to WT females (**Figure 1F**). Interestingly, young Hom females also showed increased VLDL- and intermediate-density lipoprotein (IDL)/LDL-cholesterol levels compared to the rest of the groups (**Figure 1F**). Likewise, young Het and Hom males also showed increased VLDL-triglycerides levels compared to WT males (**Figure 1G**). Finally, young Hom females exhibited elevated VLDL-triglyceride levels compared to young WT females, while adult Het and Hom females had higher levels compared to adult WT females (**Figure 1H**).

Lastly, to investigate whether mouse APOE CC alters life expectancy, we measured the probability of survival of WT, Het and Hom APOE CC mice up to 24 months of age and found a similar life expectancy between groups (**Figure 1I**).

### Minimal effects of murine APOE CC on Aβ pathology in 5xFAD mice

The human case reported by *Arboleda et al*., who carried the PSEN1 E280A and the homozygous APOE3 CC mutations showed high brain amyloid burden despite not developing dementia ever into her 70s, several decades after PSEN1 E280A carriers in her kindred typically exhibit severe memory impairments.^14^ To investigate the effects of murine APOE CC on Aβ pathology, we crossed the APOE CC mice with the Aβ-plaque forming 5xFAD mouse model, generating three different groups: 5xFAD mice heterozygous (5xFAD;Het) and homozygous (5xFAD;Hom) for the APOE CC mutation and the control mice (5xFAD;WT). In the 5xFAD model, Aβ plaques are detected by 2 months of age^30^ and become more pronounced by 4 months^31^. Notably, females exhibit more severe pathology than males.^32^ Therefore, we collected the brains from 6 month-old mice and prepared them for histological and biochemical analysis. Aβ-plaque staining with 4g8 (**Figure 2A**) revealed no significant genotype differences in the number of Aβ plaques throughout the whole brain (**Figure 2B**). However, when groups were split by sex, females exhibited increased plaque levels compared to males with 5xFAD;Hom males showing reduced pathology compared to 5xFAD;Het and a trend towards less pathology compared to 5xFAD;WT males (**Figure 2C**). ELISA analysis of Aβ levels in the PBS-soluble (**Figure 2D**) and PBS-insoluble (**Figure 2E**) cortical and hippocampal fractions showed no significant differences among genotypes or sexes. However, cortical insoluble Aβ levels showed a trend towards reduced levels in 5xFAD;Hom compared to 5xFAD;WT and 5xFAD;Het mice (**Figure 2E**).

**Figure 2.**
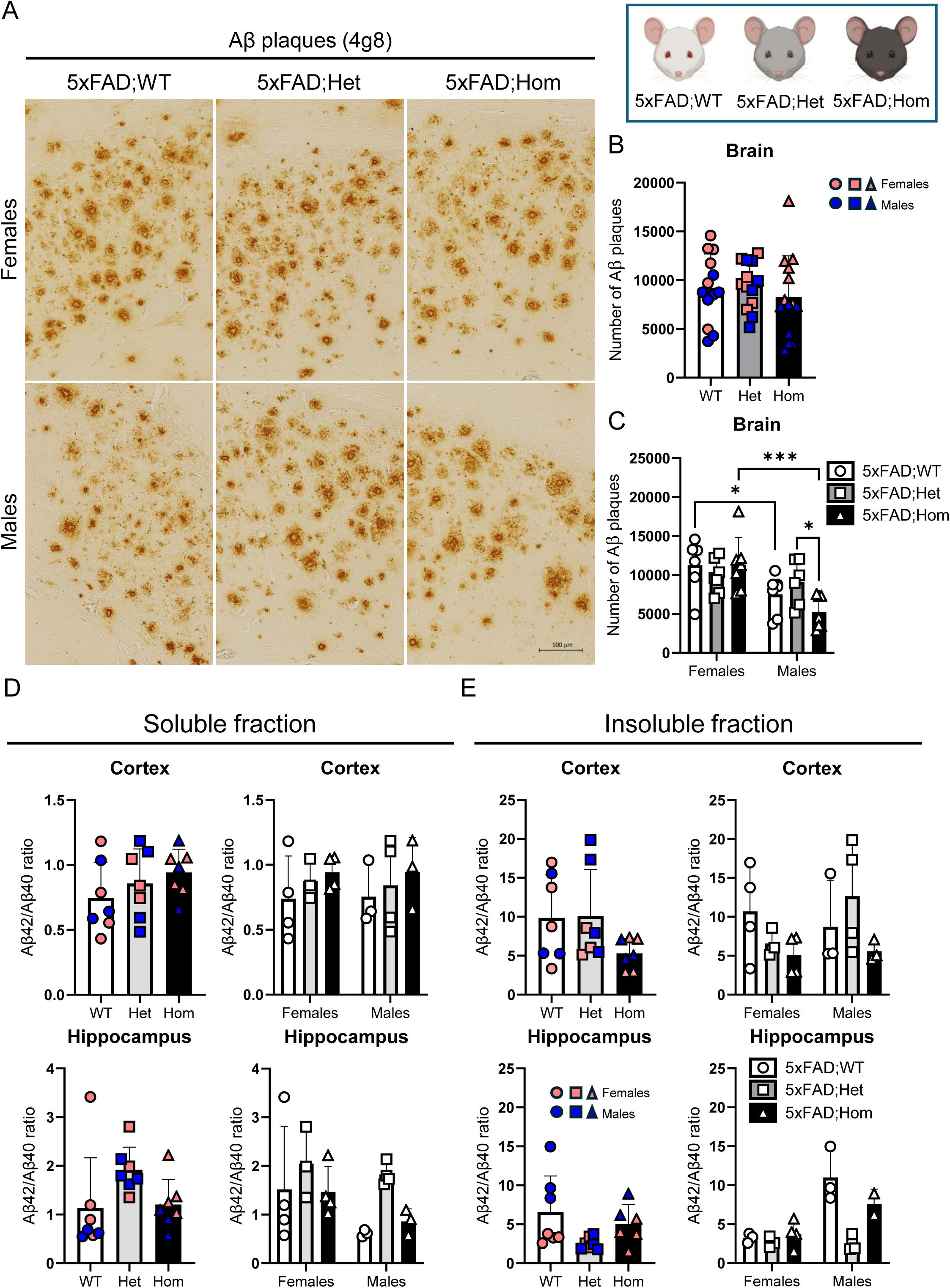
Minimal effects of APOE CC on Aβ plaque pathology. **A** Immunohistochemistry images of Aβ plaque (4g8) in the hippocampus of 6 months-old mice (scale bar = 100 µm). **B-C** Quantitative analysis of brain Aβ plaques with females and males combined (**B**, n = 13-14 per genotype) or split by sex (**C**, sex effect: F (1,34) = 15.82, *p* = 0.0003, n = 6-7 mice/group). Data represents the average number of Aβ plaques ± SD per section (6 sections/mouse). One- and two-way ANOVA followed by Tukey’s post hoc were used as statistical tests. * *p* < 0.05. **D-E** ELISA analysis of PBS-soluble (**D**) and PBS-insoluble (**E**) fractions in the cortex and the hippocampus measuring Aβ42/Aβ40 ratio. Data represents Aβ42/Aβ40 ratio levels ± SD (n = 7 mice per genotype or n = 3-4 mice/sex and genotype). One- and two-way ANOVA followed by Tukey’s post hoc was used as statistical tests.

### No effect of murine APOE CC on tau pathology in PS19 mice

Next, we examined the role of murine APOE CC on tau pathology due to mutant *MAPT* overexpression by crossing APOE CC mice with tau tangle-forming PS19 mice to generate three different groups: PS19 mice heterozygous (PS19;Het) and homozygous (PS19;Hom) for APOE CC and control mice (PS19;WT). PS19 mice show tau tangle pathology and neuronal loss by 6 months of age,^33^ with males exhibiting a more severe and progressive pathology than females.^34^ We collected brains from 8 month-old mice and prepared them for histological and biochemical analysis. We quantified the number of phosphorylated tau positive inclusions using the AT8 antibody (pSer202/Thr205 tau, **Figure 3A**) and found no significant differences between groups (**Figure 3B-C**). Likewise, biochemical analysis of PBS-soluble (**Figure 3D**) and PBS-insoluble (**Figure 3E**) cortical and hippocampal fractions showed no differences between groups, suggesting that murine APOE CC does not directly affect tau pathology formed by mutant *MAPT* overexpression.

**Figure 3.**
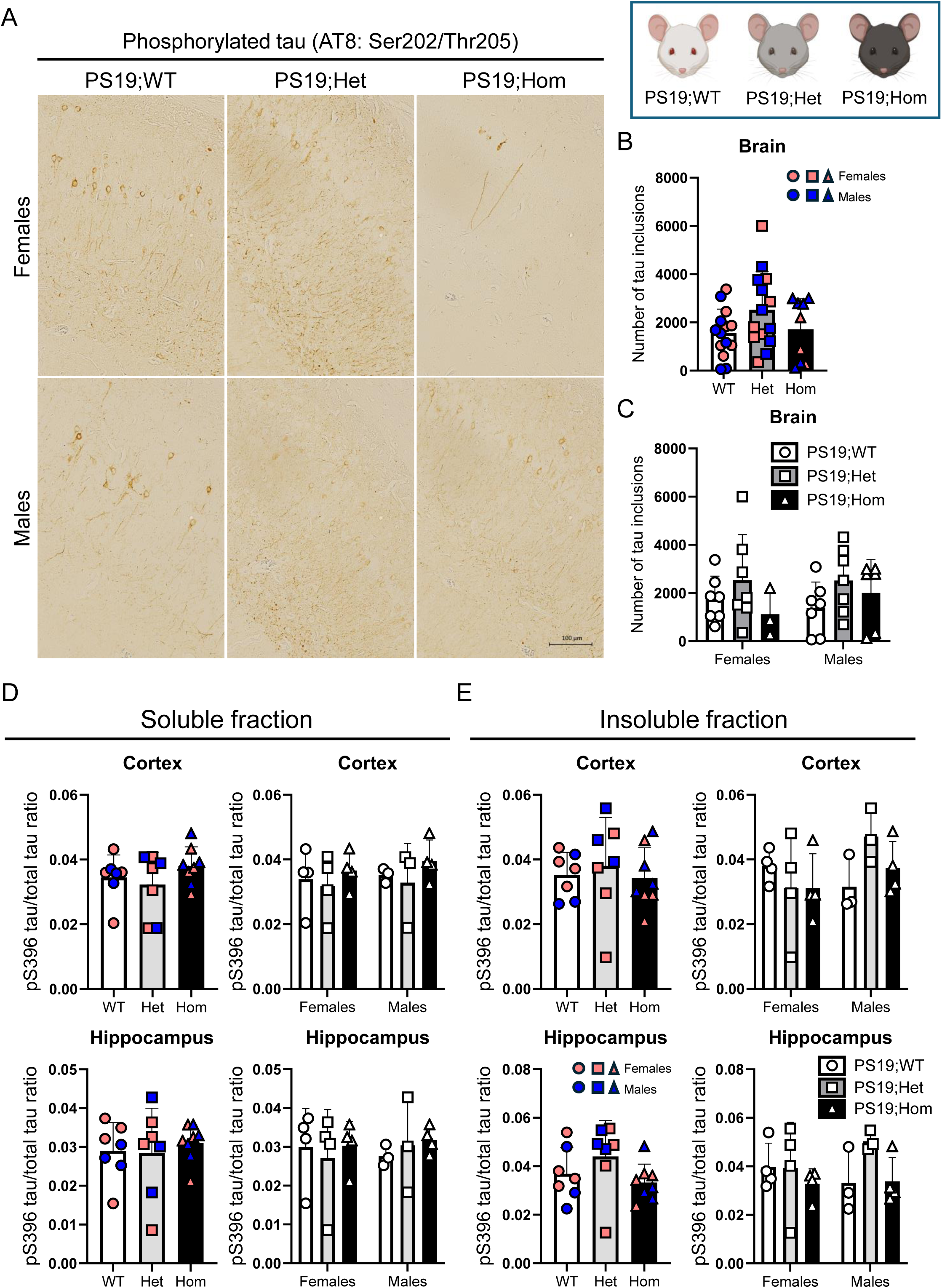
No direct effects of murine APOE CC on tau pathology in PS19 mice. **A** Immunohistochemistry images of pS202/T205 tau (AT8) in the hippocampus of 8 months-old mice (scale bar = 100 µm). **B-C** Quantitative analysis of brain number of tau positive inclusions with females and males combined (**B**, n = 9-14/genotype) or split by sex (**C**, n = 3-7 mice per group). Data represents the average inclusion number ± SD per section (6 sections/mouse). One- and two-way ANOVA followed by Tukey’s post hoc were used as statistical tests. **D-E** ELISA analysis of PBS-soluble (**D**) and PBS-insoluble (**E**) fractions in the cortex and the hippocampus measuring pS396/total tau ratio. Data represents pS396/total tau ratio levels ± SD (n = 7 mice/genotype or n = 3-4 mice/sex and genotype). One- and two-way ANOVA followed by Tukey’s post hoc was used as statistical tests.

### Murine APOE CC reduces tau pathology formed near Aβ plaques in female 5xFAD mice after intracerebral PHF-tau injections

To test whether murine APOE CC influences tau pathology that forms near Aβ plaques after injection of aggregated tau into the brain, we injected PHF-tau isolated from human AD brain, as previously described,^35,36^ into 4-month-old 5xFAD;WT and 5xFAD;Hom mice. 3 months post injection (mpi), brains were collected and processed for histological analysis. Similar to above, Aβ plaque pathology assayed by 4g8 staining across the brain (**Figure 4A**) revealed no significant differences between genotypes (**Figure 4B**). However, when analyzed by sex, female mice showed elevated numbers of Aβ plaques compared to male mice (**Figure 4C**). Interestingly, only 5xFAD;WT females had increased number of Aβ plaques than 5xFAD;WT males, while this differences was not significant between 5xFAD;Hom females and 5xFAD;Hom males (**Figure 4C**), suggesting that APOE CC can reduce differences between the sexes in Aβ plaque burden in 7-month-old 5xFAD animals in the context of PHF-tau injection. Analysis of tau pathology with AT8 staining throughout the brain (**Figure 4D**) revealed a trend to reduce the number of tau inclusions in 5xFAD;Hom compared to 5xFAD;WT mice (**Figure 4E**). However, when the data were split by sex, there was a sex effect with 5xFAD;WT females showing higher amount of pathology than 5xFAD;WT males, a genotype effect with 5xFAD;Hom females showing reduced number of tau inclusions than 5xFAD;WT females, and a sex x genotype interaction showing that the APOE CC variant reduces tau pathology in 5xFAD females but not in males (**Figure 4F**). Interestingly, there was a significant positive correlation between the number of Aβ plaques and tau inclusions (**Figure 4G**), suggesting that Aβ pathology influences tau pathology levels.

**Figure 4.**
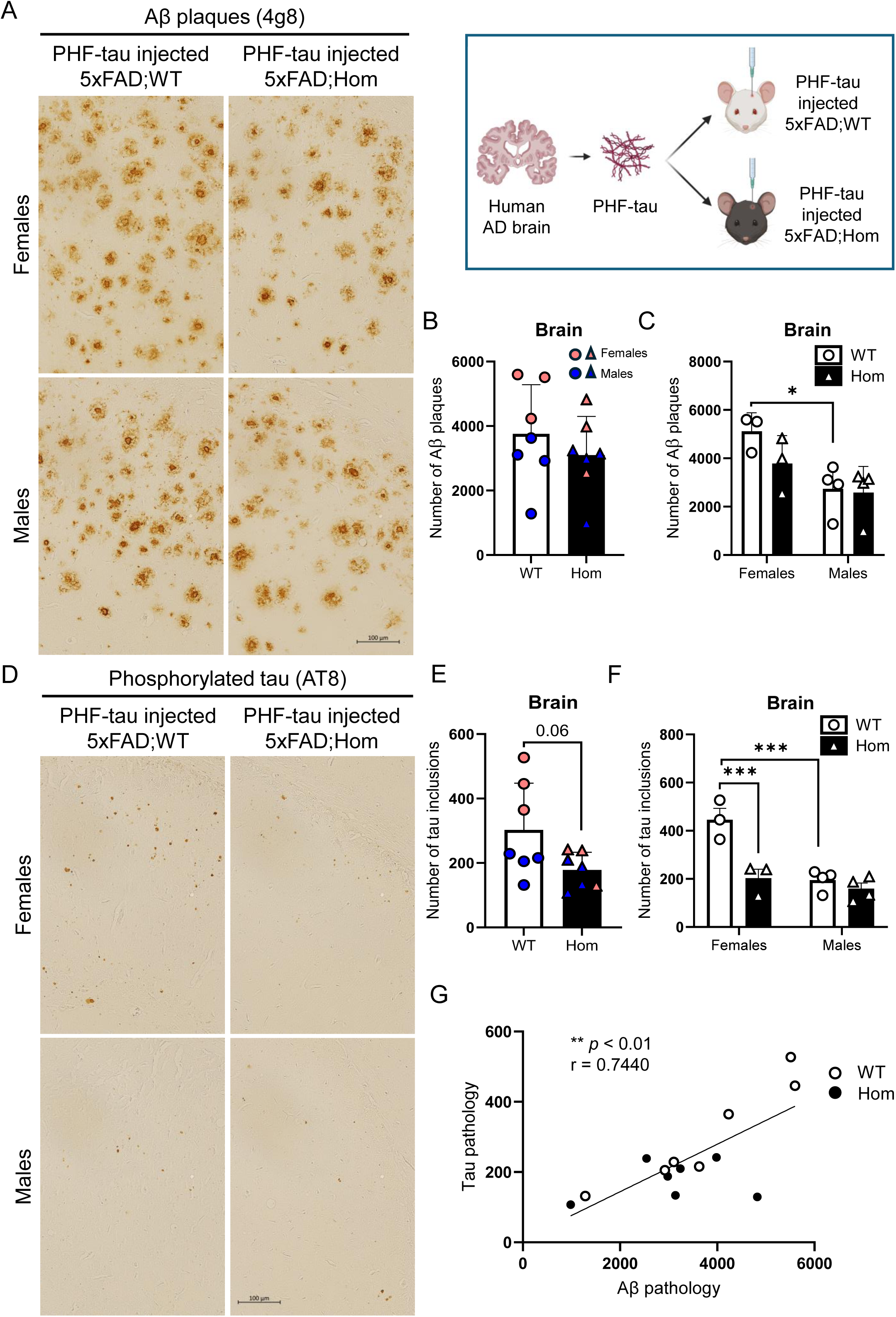
APOE CC mutation reduces Aβ plaque and Aβ-induced tau pathologies in PHF-tau injected 5xFAD mice. **A** Immunohistochemistry images of Aβ plaque (4g8) in the hippocampus of 7 months-old injected 5xFAD mice (scale bar = 100 µm). **B-C** Quantitative analysis of brain Aβ plaque with females and males combined (**B**, n = 7-8 mice per genotype) or split by sex (**C**, sex effect: F (1,10) = 10.58, *p* = 0.0087, n = 3-4 mice/group). Data represents the average Aβ plaque number ± SD per section (15 sections/mouse). Unpaired t-test and two-way ANOVA followed by Tukey’s post hoc were used as statistical tests. * *p* < 0.05. **D** Immunohistochemistry images of pS202/T205 tau (AT8) in the hippocampus of 7 months-old injected 5xFAD mice (scale bar = 100 µm). **E-F** Quantitative analysis brain tau inclusions with females and males combined (**E**, n = 7-8 mice per genotype) or split by sex (**F**, sex effect: F (1,10) = 21.91, *p* = 0.0009; genotype effect: F (1,10) = 19.60, *p* = 0.0013; sex x genotype interaction: (F (1,10) = 10.83, *p* = 0.0081), n = 3-4 mice/group). Data represents the average tau inclusion number= ± SD per section (15 sections/mouse). Unpaired t-test and two-way ANOVA followed by Tukey’s post hoc were used as statistical tests. *** *p* < 0.001. **G** Correlation analysis between Aβ and tau pathological levels in injected 5xFAD mice (*p* < 0.01, R^2^ = 0.5536, r = 0.7440) analyzed using two-tailed Pearson’s correlation coefficient (r). ** *p* < 0.01.

### Strong sex, but subtle genotype effects on behavior in APOE CC mice crossed to 5xFAD, PS19, or PHF-tau injected 5xFAD mice

To examine the effects of APOE CC on behaviors that involve motor function, anxiety and cognition in the different Aβ and/or tau models described above, we performed a battery of behavioral tests including nest building (NB), elevated zero maze (EZM), open field (OF), novel object recognition (NOR) and contextual and cued fear conditioning (FC).

In 5xFAD mice, there were sex significant differences in NB, but no genotype effects due to APOE CC expression. In the OF, there were also sex-dependent effects on the distance moved with 5xFAD;WT and 5xFAD;Het females showing more activity than males of the same genotype, but no differences between 5xFAD;Hom females and males. There was also an effect of sex on the time spent in the center of the arena, with males spending more time than females, without APOE CC genotype effects. Finally, during the NOR, there was an effect of sex and genotype on the activity levels, with higher distance moved in females than males and in 5xFAD;Het compared to 5xFAD;WT and 5xFAD:Hom (**Figure 5A**).

**Figure 5.**
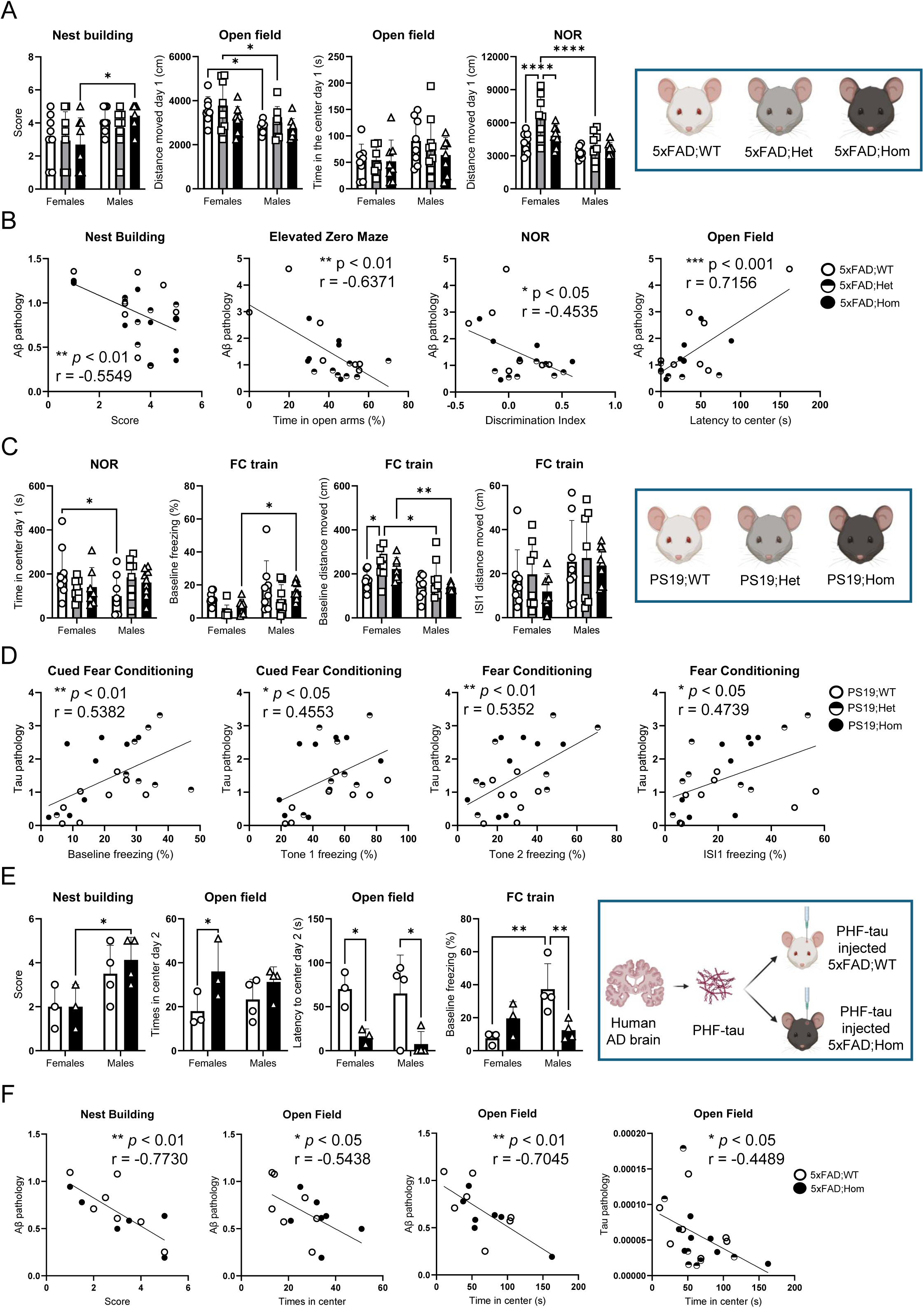
APOE CC variant has subtle effects on the behavior of 5xFAD, PS19 and PHF-tau injected 5xFAD mice. **A** Behavioral NB (sex effect: F (1,42) = 9.179, *p* = 0.0042), OF (distance moved, sex effect: F (1,42) = 9.967, *p* = 0.0029; time in the center, sex effect: F (1,42) = 4.986, *p* = 0.0309) and NOR (sex effect: F (1,42) = 21.99, *p* <0.0001; genotype effect: F (2,42) = 6.844, *p* = 0.0027) tests in 6 months-old 5xFAD;WT, 5xFAD;Het and 5xFAD;Hom (n = 8 mice/group). Two-way ANOVA followed by Tukey’s post hoc was used as a statistical test. **B** Correlation analysis between behavior and Aβ pathological levels was performed using two-tailed Pearson’s correlation coefficient (r) (NB: *p* < 0.01, R^2^ = 0.3080, r = −0.5549; EZM: *p* < 0.01, R^2^ = 0.4058, r = −0.6371; NOR: *p* < 0.05, R^2^ = 0.2057, r = −0.4535; OF: *p* < 0.001, R^2^ = 0.5121, r = 0.7156). * *p* < 0.05, ** *p* < 0.01, *** *p* < 0.001. **C** Behavioral NOR (sex x genotype interaction: F (2,42) = 3.978, *p* = 0.0262) and FC train (baseline freezing: sex effect: F (1,42) = 11.05, *p* = 0.0019; genotype effect: F (2,42) = 3.743, *p* = 0.0319; baseline distance moved: sex effect: F (1,42) = 13.51, *p* = 0.0007; genotype effect: F (2,42) = 3.753, *p* = 0.0317; ISI1, sex effect: F (1,42) = 4.573, *p* = 0.0383) in 8 months PS19;WT, PS19;Het and PS19;Hom (n = 8 mice/group). Two-way ANOVA followed by Tukey’s post hoc was used as a statistical test. * *p* < 0.05, ** *p* < 0.01. **D** Correlation analysis between behavior and tau pathological levels was done using two-tailed Pearson’s correlation coefficient (r) (baseline freezing: *p* < 0.01, R^2^ = 0.2896, r = 0.5382; tone 1 freezing: *p* < 0.05, R^2^ = 0.2073, r = 0.4553; baseline distance moved: *p* < 0.01, R^2^ = 0.2865, r = 0.5352; ISI1 distance moved: *p* < 0.05, R^2^ = 0.2246, r = 0.4739). * *p* < 0.05, ** *p* < 0.01. **E** Behavioral NB (sex effect: F (1,10) = 9.242, *p* = 0.0125), OF (Times in center, genotype effect: F (1, 10) = 6.573, *p* = 0.0282; latency to center, genotype effect: F (1, 10) = 14.57, *p* = 0.0034) and CFC (sex x genotype effect: F (1,10) = 11.06, *p* = 0.0077) in 7 months-old PHF-tau injected 5xFAD;WT and 5xFAD;Hom mice (n = 3-4 mice/group). Two-way ANOVA followed by Tukey’s post hoc was used as a statistical test. **F** Correlation analysis between behavior and Aβ/tau pathological levels was done using two-tailed Pearson’s correlation coefficient (r) (NB: *p* < 0.01, R^2^ = 0.5975, r = −0.7730; OF: frequency in the center: *p* < 0.05, R^2^ = 0.2957, r =-0.5438; time spent in the center: *p* < 0.01, R^2^ = 0.4964, r = −0.7045; tau pathology/time spent in the center: *p* < 0.01, R^2^ = 0.2016, r = −0.4489. Data represents the score, distance moved (cm), time in or latency to the center (s) and percentage of freezing ± SD. * *p* < 0.05, ** *p* < 0.01, *** *p* < 0.001, **** *p* < 0.0001.

To investigate whether the level of pathology in each mouse correlates with its behavioral performance, we performed a correlation analysis. Interestingly, there were significant negative correlations between Aβ pathology and NB score, time spent in the open areas during EZM, and discrimination index during NOR, and a significant positive correlation with the latency to go to the center during OF (**Figure 5B**).

In PS19 mice, there was a sex x genotype interaction in the time spent in the center during NOR, with PS19;WT females spending more time than PS19;WT males, and male PS19;Het and PS19;Hom spending more time than females of the same genotypes. In the CFC, there were sex and genotype effects on the baseline freezing levels, with males showing higher freezing levels than females, especially in PS19;Hom, and with PS19;WT mice showing higher levels than PS19;Het and PS19;Hom mice. Similarly, there were also sex and genotype effects on the baseline activity levels during the contextual FC, with PS19;Het and PS19;Hom females showing higher levels than PS19;Het and PS19;Hom males, respectively, and female PS19;Het being more active than PS19;WT females. Finally, there was an effect of sex on the freezing levels during the inter-stimulus interval (ISI) 1, with males showing elevated levels than females (**Figure 5C**).

Similar to above, we decided to perform a correlation analysis of the level of pathology in each mouse with its behavior. Notably, there were significant positive correlations between the amount of tau pathology and freezing and during baseline and tone 1 and the distance moved during the baseline and ISI1 during the CFC (**Figure 5D**).

Lastly, we analyzed the behavioral effects of APOE CC in PHF-tau injected 5xFAD mice. We found an effect of sex on the NB score, with males showing a better score than females. During OF, there was a genotype effect on the frequency and latency to the center, with PHF-tau 5xFAD;Hom exploring more times and earlier the center than 5xFAD;WT mice. Finally, there was a sex x genotype interaction on the baseline freezing levels during the CFC, with PHF-tau injected 5xFAD;WT males showing higher levels than 5xFAD;WT females and 5xFAD;Hom males showing decreased levels than 5xFAD;WT males (**Figure 5E**).

We also performed a correlation analysis between the levels of Aβ/tau pathologies and the behavioral performance in each mouse individually. Interestingly, there were significant negative correlations between the amount of Aβ pathology and NB score, frequency in the center and time spent in the center during OF, which also correlate with the amount of tau pathology (**Figure 5F**).

The rest of the data from motor, anxiety and cognitive tests can be found in **Supplementary Figures 1-3**.

### APOE CC mice showed reduced α-syn pathology after intracortical α-syn PFF injection

To examine the effects of murine APOE CC on LB pathology, we performed intracortical α-syn PFF injections in 4-month-old Het and Hom APOE CC and control WT littermate mice. 4 mpi brains were collected and processed for histological analysis. Interestingly, using a specific antibody against phosphorylated α-syn (pS129, EP1536Y, **Figure 6A**), we found a genotype effect with Het and Hom mice showing reduced α-syn inclusion number in the ipsilateral hemisphere compared to WT mice (**Figure 6B**). To analyze the spread of α-syn, we measured the number of α-syn inclusions in the hemisphere contralateral to the injection site and found a genotype effect with WT mice showing more α-syn inclusions compared to Hom, and a trend to more α-syn inclusions compared to Het mice (**Figure 6C**). When the α-syn inclusion number in both hemispheres was quantified together, we found a genotype effect with Het and Hom mice showing reduced α-syn inclusion number compared to WT mice (**Figure 6D**).

**Figure 6.**
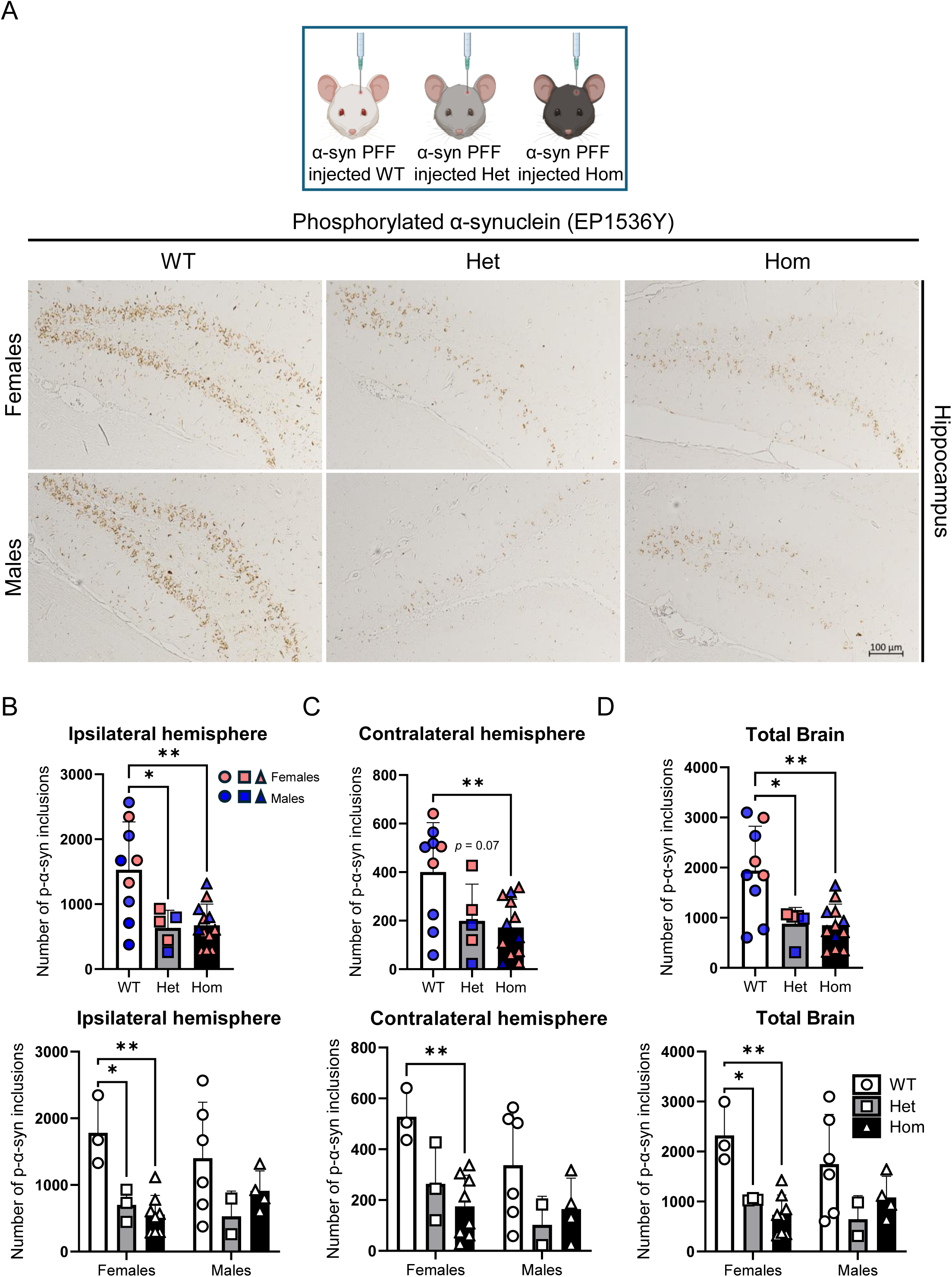
APOE CC expression reduces α-syn pathology after intracortical α-syn PFF injection. **A** Immunohistochemistry images of pS129 α-syn (EP1536Y) in the hippocampus of 8 months-old mice 4 mpi (scale bar = 100 µm). **B-D** Quantitative analysis of α-syn inclusion number in the injected (**B**, genotype effect: F (2,23) = 8.749, *p* = 0.0015), non-injected (**C**, genotype effect: F (2,23) = 0.5.815, *p* = 0.009) and both (**D**, genotype effect: F (2,23) = 9.126, p = 0.0012) hemispheres with females and males combined (n = 5-12 mice/genotype) or split by sex (n = 2-6 mice/group). Data represents the average α-syn inclusion number ± SD per section (15 sections/mouse). One- and two-way ANOVA followed by Tukey’s post hoc were used as statistical tests. * *p* < 0.05, ** *p* < 0.01.

### APOE CC mice showed reduced spinal cord pathology after hindlimb gastrocnemius α-syn PFF injection

To explore the effects of APOE CC on α-syn spread from the periphery into the CNS, we crossed the APOE CC mice with the A53T mouse model expressing this familial PD α-syn mutation.^37^ This model is characterized by accelerated LB pathology induction and cell death after intracortical α-syn PFF injection, and a retrograde trans-synaptic spread of LB pathology from the periphery to the CNS after hindlimb gastrocnemius muscle α-syn PFF injection, spreading up the motor system and reaching the mesencephalon 4 mpi and the motor cortex 8 mpi.^37^ Therefore, we generated heterozygous (A53T;Het) and homozygous (A53T;Hom) APOE CC mice, all with heterozygous A53T expression, and the control mice (A53T;WT), and performed muscle α-syn PFF injection at 4 months old. Half of the mice were euthanized 4 mpi, while the other half was euthanized 8 mpi. Brains and spinal cords were collected and processed for histological approaches using an antibody against phosphorylated α-syn (**Figure 7A-B**). In the 4 mpi group, there was an effect of sex in the number of α-syn inclusions in the mesencephalon, with female mice showing increased number than males and a tendency to have less inclusions in A53T;Hom compared to A53T;WT mice (**Figure 7C**). In the spinal cord, there was also a sex effect, with females showing an elevated number of α-syn inclusions than males in A53T;WT and A53T;Het mice, but not in A53T;Hom mice and with A53T;WT females trending to have more α-syn inclusions than A53T;Hom females (**Figure 7D**). In the 8 mpi group, the number of α-syn inclusions in the mesencephalon and the spinal cord were higher than in the 4 mpi group. In the mesencephalon, there was an effect of sex with increased number of α-syn inclusions in females than males, but not significant differences between genotypes (**Figure 7E**). In the spinal cord, there was a sex effect with increased number of α-syn inclusions in females than males and a genotype effect with A53T;WT females showing elevated number of α-syn inclusions than A53T;Het and A53T;Hom females (**Figure 7F**).

**Figure 7.**
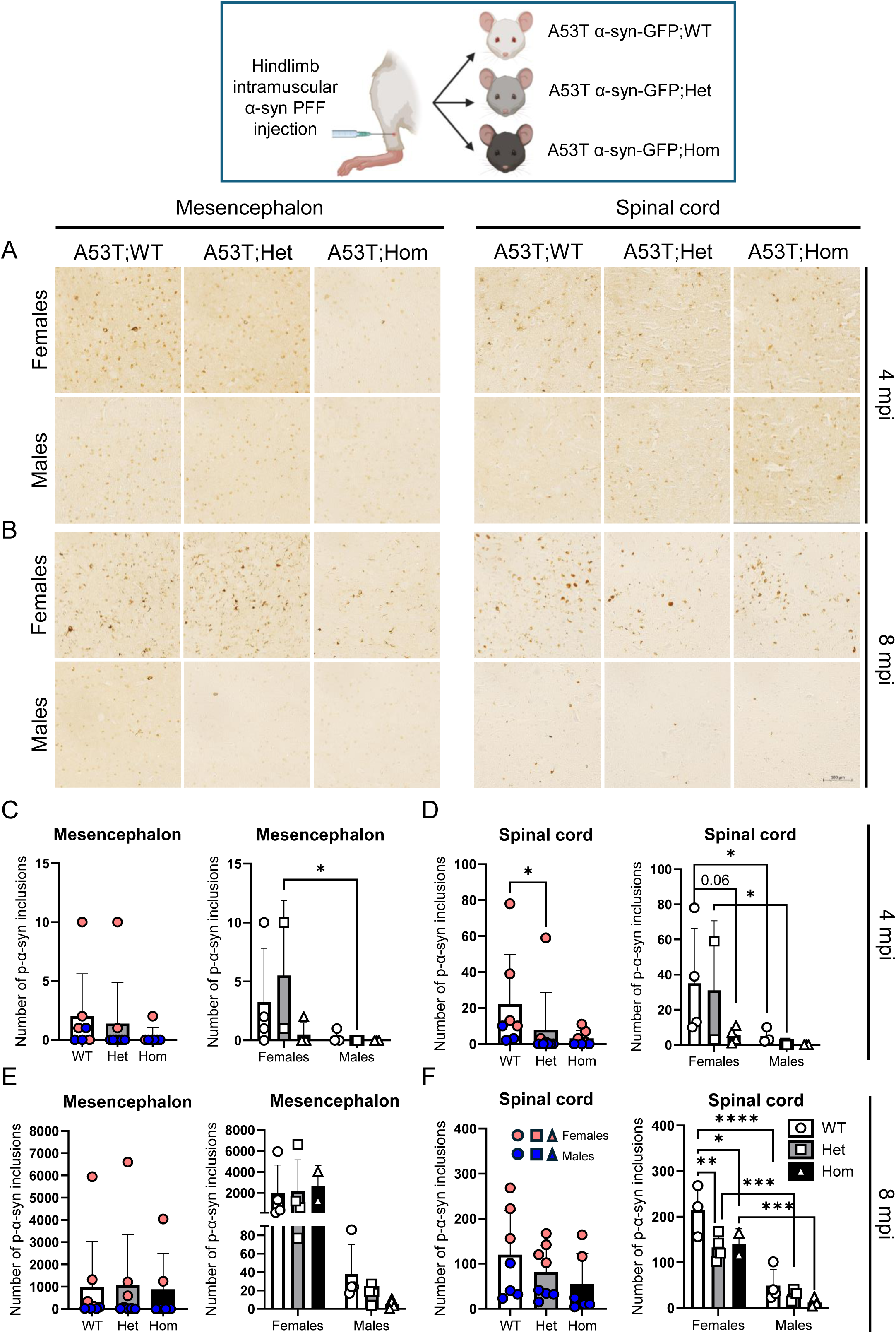
APOE CC expression reduces α-syn pathology in the spinal cord of female A53T mice after intramuscular α-syn PFF injection. **A-B** Immunohistochemistry images of pS129 α-syn (EP1536Y) in the mesencephalon and spinal cord of 4 (**A**) and 8 (**B**) mpi A53T mice injected with α-syn PFF in the hindlimb gastrocnemius muscle at 4 months-old (scale bar = 100 µm). **C-F** Quantitative analysis of total α-syn inclusion number in the mesencephalon (**C**, sex effect: F (1,16) = 6.490, *p* =0.0215. **E**, sex effect: F (1,16) = 7.512, *p* = 0.0145) and the spinal cord (**D**, sex effect: F (1,16) = 8.276, *p* = 0.0110. **F**, sex effect: F (1,15) = 90.70, *p* < 0.0001; genotype effect: F (2,15) = 6.725, *p* = 0.0082) of injected mice with females and males combined (n = 7-8 mice/genotype) or split by sex (n = 2-6 mice/group). Data represents total number of α-syn positive inclusions ± SD (15 sections/mouse). One- and two-way ANOVA followed by Tukey’s post hoc were used as statistical tests, except for α-syn inclusions in the spinal cord in the 4 mpi, in which the SD among groups were different and Brown-Forsythe ANOVA and Welch’s ANOVA tests were used. * *p* < 0.05, ** *p* < 0.01, *** *p* < 0.001.

## Discussion

Arboleda-Velasquez et al. described a case of a carrier of the normally fully penetrant familial AD PSEN1-E280A mutation, but homozygous for the *APOE3 CC* variant, who remarkably showed resistance to tau pathology in certain brain regions and dementia development.^14^ PSEN1-E280A carriers of the same kindred develop mild cognitive impairments and dementia at 45- and 50-year-old, respectively.^38^ However, this homozygous *APOE3 CC* patient only developed subtle short-term memory impairments in her 70s, ∼30 years later than dementia was expected. This suggested a protective effect against AD mediated by *APOE CC* and raised hope for potential therapeutic strategies based on this new result. Intriguingly, APOE has also been implicated in the biology of LB disorders like PD and DLB, although before our work, there has been no data that we are aware of about the possible ability of the APOE CC variant being protective in LB disorders. The low allele frequency of *APOE CC* in the human population makes it hard to investigate the potential protective mechanisms implicated in neurodegeneration related to Aβ, tau or α-syn, and the creation of model systems is important for answering these questions. Here, we developed a gene-targeted mouse expressing murine APOE R128S CC and used histological, biochemical and behavioral approaches to examine the effects of murine APOE CC on Aβ plaque, tau tangle and LB pathologies. We found protective effects of murine APOE CC on Aβ-induced tau pathology in 5xFAD mice after the injection of PHF-tau, but not direct effects on Aβ plaque in 5xFAD mice or tau pathology burden in PS19 mice. We also found, for the first time, protective effects of murine APOE CC on LB with reduction of α-syn pathology after intracortical and intramuscular α-syn PFF injection in non-transgenic and A53T mice, respectively.

After the Arboleda-Velasquez et al. report, several studies have explored the role of APOE CC in AD models using humanized APOE^18,19,25^ and single-point mutant APOE mice,^26^ adeno-associated viral APOE vectors^20,24^ and in vitro strategies (**Supplementary Table 1**).^21,23^ Specifically, Nelson et al. knocked in the human *APOE4 CC* allele and crossed these mice to PS19 and detected a reduction in tau pathology, hippocampal atrophy and in the levels of GFAP, S100β, Iba1 and CD68, when APOE4 CC was homozygous, and similar results (except no reduction in tau pathology) in heterozygous APOE4 CC mice.^18^ Chen et al. knocked in human *APOE3 CC* and found higher levels of plasma cholesterol and a reduction in the total number of Aβ plaques and plaque-associated tau pathology after the intracerebral injection of PHF-tau in APP/PSEN1 female mice. However, there was no effect on tau pathology levels in the absence of Aβ pathology and APOE3 CC was not able to rescue memory impairments on the Y maze or fear conditioning.^19^ Naguib et al. showed a reduction in tau pathology and decreased synaptic and myelin loss in PS19 mice with human *APOE3 CC* knocked in.^25^ Tran et al. used a strategy more similar to ours to create a single-point mutant mouse expressing murine APOE CC (R128S) and found increased plasma cholesterol and reduced Aβ plaque load when crossed with 5xFAD mice, but no effects in tau pathology when crossed with PS19 mice. They also detected no changes in hindlimb clasping score in the APOE CC/PS19 mice and no changes in the time spent in the open arms in the elevated plus maze or the center during the OF in APOE CC/PS19 and APOE CC/5xFAD mice.^26^

Other groups have used an adeno-associated viral injection strategy to express human APOE CC. Gunaydin et al. reported reduced Aβ and tau pathologies in APP/PSEN1/TRE4 and PS19/TRE4 mice, respectively, and behavioral rescue in NB, Y maze, NOR and Barnes maze tests after hippocampal adeno-associated viral injection of APOE2 CC.^20^ Chen et al. used adeno-associated viral injection of APOE3 CC into the Locus Coeruleus and found reduced tau pathology in PS19 mice and reduced Aβ and Aβ-induced tau pathology after the injection of Tau-PFFs in 5xFAD. They also found improved learning and memory in the Morris water maze and fear conditioning in 5xFAD and PS19 mice.^24^

In addition, several studies have explored the mechanism of human APOE CC’s protective role in vitro. Murakami et al. found that APOE3 CC astrocytes in culture limit tau propagation within admixed PSEN1 mutant neurons^21^ and Sun et al. showed that APOE3 CC microglia are resistant to Aβ42-induced lipid peroxidation, ferroptosis and impaired phagocytosis, and reduce tau pathology within PSEN1 mutant neurons,^23^ highlighting cell type-specific effects of APOE CC.

In agreement with several of these in vivo mouse model studies,^19,26^ we have also observed increased cholesterol levels in the plasma of young and adult Het and Hom APOE CC mice. However, in contrast to Tran et al., we also found increased plasma triglyceride levels as it seen in humans with APOE CC. This discrepancy may be due to the different experimental approaches used, as Tran et al. used an enzymatic colorimetric assay measuring bulk-concentration, combining male and female plasma samples, and we performed size-based chromatography, measuring lipoprotein distribution in males and females independently. In contrast to the effects seen using human APOE CC in humanized mice^18,25^ or adeno-associated viral vectors,^20,24^ and in agreement with Tran et al., we did not see any direct effect of the murine APOE CC on the total number of tau inclusions in PS19 mice. Chen et al. 2024 reported a protective effect of human APOE3 CC against tau pathology in APP/PSEN1 female mice after the injection of human AD-tau,^19^ an effect that was also seen by Chen et al. 2025 using recombinant tau PFF injections in 5xFAD mice with adeno-associated human APOE3 CC expression.^24^ Interestingly, we also detected a significant reduction in Aβ-induced tau pathology after the injection of PHF-tau purified from AD brains mediated by murine APOE CC in 5xFAD female mice. Although the number of tau aggregates significantly correlated with the number of Aβ plaques, the effects on tau pathology were stronger, suggesting that murine APOE CC might not only have direct effects on Aβ pathology but also on the ability of Aβ to promote tau pathology formation. Although AD mouse models have been shown to exhibit sex-differences in pathology and behavior,^34,39,40,41,42^ none of the previous studies exploring the effects of APOE CC on AD-like pathology and behavior measured sex as a biological variable.^18,19,20,24,25,26^ Importantly, our study is the only one that reported sex differences in the effects of APOE CC against AD-like pathology.

To explore the effects of murine APOE CC on the behavior, we performed a battery of behavioral tests in PS19 and 5xFAD mice to look for changes in motor and cognitive function and anxiety mediated by murine APOE CC in a sex-dependent manner. This includes the first tests we are aware of in 5xFAD mice homozygous for APOE CC and injected with PHF-tau (to seed Aβ-induced tau pathology). In non-injected APOE CC/5xFAD mice, we found strong sex-differences in behavior, with females showing lower nest building scores and a more anxiety-like phenotypes than males. However, 5xFAD;Hom females exhibit reduced activity levels during the OF (similar to males), suggesting that homozygous APOE CC reduces this anxiety phenotype in 5xFAD females. In PS19 mice, although we found no changes in tau pathology, we found a sex x genotype interaction in the time spent in the center during NOR, showing that both heterozygous and homozygous APOE CC expression reduced anxiety in PS19 males but increased anxiety in PS19 females. In FC, we also found sex effects on the basal freezing levels with males showing higher levels than females, but more interestingly, PS19;WT mice showed higher levels than PS19;Het and PS19;Hom mice, suggesting that both heterozygous and homozygous APOE CC genotypes reduce anxiety in PS19 mice. Finally, in the PHF-tau injected 5xFAD mice, there was also a genotype effect in the frequency and latency to the center with Hom mice showing lower anxiety levels than WT mice. As expected, in all the different cohorts, behavioral performance and anxiety correlated significantly with the levels of Aβ and/or tau pathologies, suggesting that the amount of pathology influences these behavioral responses.

Although the effects of APOE in PD have been less studied than in AD, *APOE* genotype also correlates with PD pathology progression and dementia risk, and *APOE4* carriers show elevated pathological α-syn levels.^5,6^ However, to our knowledge, there has been no published data that we are aware of about the role of APOE CC in α-syn pathology. Therefore, we investigated the effects of murine APOE CC on LB pathology development and spread by performing intracortical α-syn PFF injections in heterozygous, homozygous APOE CC and their respective control mice. Interestingly, we found a strong reduction in the number of α-syn inclusions in both ipsilateral and contralateral brain hemispheres of heterozygous and homozygous APOE CC mice compared to WT mice. Finally, we performed intramuscular hindlimb gastrocnemius α-syn PFF injections in A53T mice to explore the role of APOE CC in LB pathology spread from the periphery to the CNS. Interestingly, A53T;Het and, especially, A53T;Hom mice showed decreased number of α-syn inclusions in the spinal cord compared to A53T;WT mice, while no differences were found in the brain, suggesting that APOE CC limits the development of α-syn pathology after peripheral α-syn PFF injection, but not its spread to the brain in this system. These results suggest that, similar to its role in reducing Aβ-induced tau pathology, murine APOE CC protects against α-syn pathology and heterozygosity is sufficient for these protective effects against LB pathology.

Although different mechanisms have been implicated in APOE CC’s protective effects,^14,18,19,20,21,22,23,24,25,27,28^ they are not mutually exclusive. Specifically, APOE CC has been shown to reduce APOE binding to the LDL receptor and HSPGs,^27,43,44^ interfering with tau uptake/spread and limiting tau pathology development.^14,18,19^ HSPGs are cell surface proteins that carry HS glycosaminoglycan chains that are negatively charged and interact electrostatically with other positively charged proteins, like tau and α-syn’s N-terminus. Interestingly, it has been proposed that HSPG are also critical mediators in the transmission of α-syn and tau, as highly sulfated domains of HSPG are key for α-syn and tau binding affinity and interaction,^45,46^ HSPG are required for α-syn and tau fibril uptake^47^ and HS mimetics block fibril uptake.^48^. Pursuing how interfering with APOE binding to HSPGs may alter α-syn and tau aggregation in the context of the CC mutation, in addition to other possible neuroprotective mechanisms, will be important in the future to understand how this single point mutation is protective and to harness this knowledge to create new therapies.

In summary, in this report we used histological, biochemical and behavioral approaches to test the effects of murine APOE CC on Aβ, tau, Aβ-induced tau and α-syn pathologies by crossing a newly generated gene-targeted murine APOE CC mouse with models of amyloidosis (5xFAD), tauopathy (PS19) and mutated α-syn overexpression (A53T). We found that murine APOE CC reduces Aβ-induced tau and α-syn pathologies in mice. Thus, these results highlight the protective function of APOE CC against different proteinopathies and opens up new avenues for studying and developing therapeutic strategies for disorders such as AD, PD and DLB.

## Data availability

Supplementary materials were available as the online version.

## Abbreviations

α-syn: α-synuclein
Aβ: Amyloid β
AD: Alzheimer’s disease
APOE: Apolipoprotein
E CC: Christchurch
CNS: Central nervous system
DAB: 3,3’-Diaminobenzidine
DLB: Dementia with Lewy body
ELISA: Enzyme linked-immuno-sorbent assay
EZM: Elevated zero maze
FC: Fear conditioning
Het: Heterozygous
HDL: High-density lipoprotein
Hom: Homozygous
HSPG: Heparan sulfate proteoglycans
IDL: Intermediate-density lipoprotein
ISI: Inter-stimulus interval
KI: Knock in
LB: Lewy body
LDL: Low-density lipoprotein
NB: Nest building
NOR: Novel object recognition
OF: Open field
PD: Parkinson’s disease
PFF: Preformed fibrils
PHF: Paired helical filaments
SD: Standard deviation
VLDL: Very low-density lipoprotein
WT: Wild type
FPLC: Fast protein liquid chromatography

## Acknowledgements

All imaging experiments could not have been possible without help from S. K. Petrie, F. Kelly, B. Jenkins and H. Bronstein in the OHSU Advanced Light Microscopy Core (RRID: SCR_009961).

## Star Methods

### Key resource table

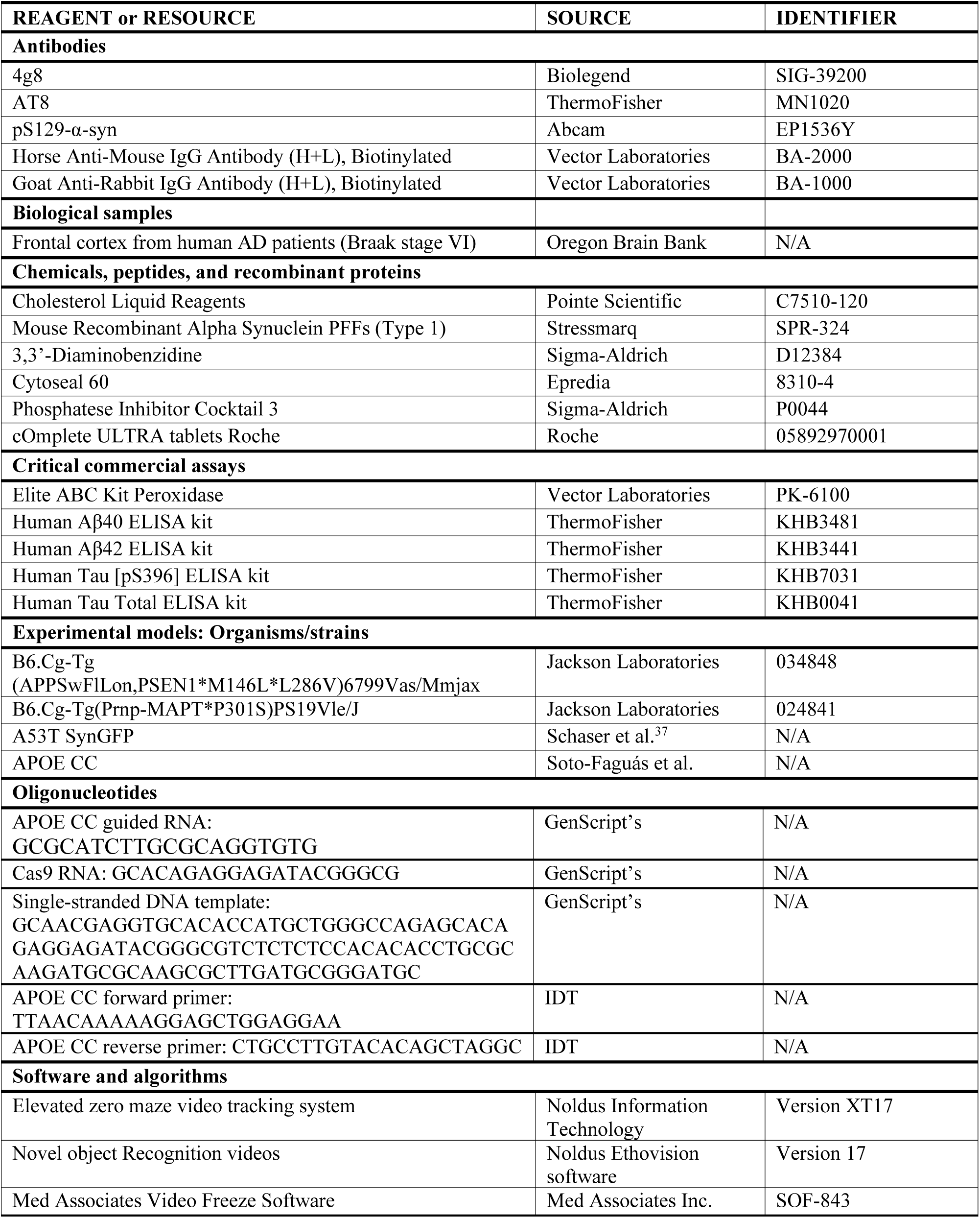
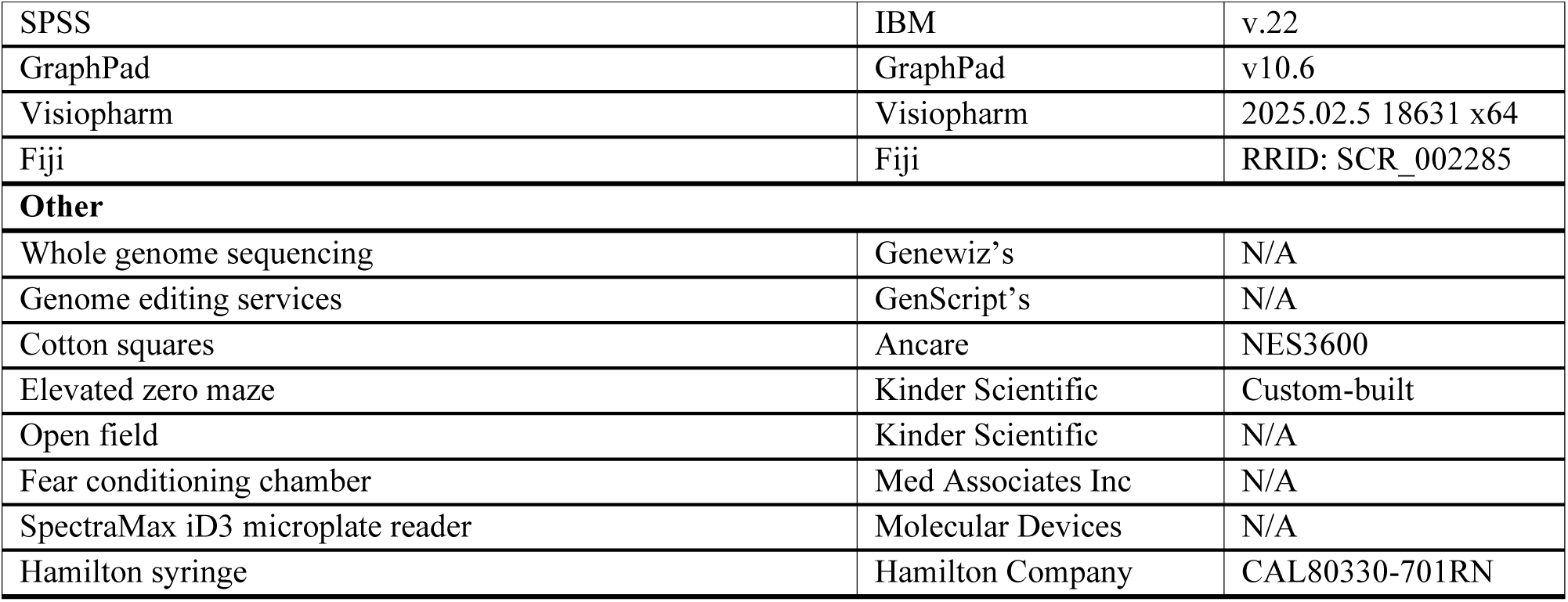

### Experimental model and study participant details

#### Animals

Animals were housed by Oregon Health & Science University (OHSU)’s Department of Comparative Medicine in a light–dark cycle, temperature and humidity-controlled vivarium, and maintained under ad libitum food and water diet. Cages were changed every 2 weeks with a maximum of 5 animals per cage. All experiments were approved by the OHSU IACUC. All experiments were performed in accordance with the relevant guidelines and regulations, and every effort was made to minimize the number of animals used and their suffering.

#### Mouse model generation

We used the OHSU Transgenic Mouse Model Core to create a founder line of the R128S APOE mice (APOE CC). We used CRISPR/Cas9-based recombination strategy with a single-stranded DNA template to introduce a CGG to TCT missense change in mouse ApoE to generate a substitution of an arginine to a serine at position 128 (R128S, **Figure 1A**). With the consultation and genome editing services of GenScript’s (Piscataway, NJ), we produced a guided RNA 10 bp downstream from the targeted codon on the reverse strand, Cas9 RNA and a single-stranded DNA template coding for the R128S mutation, for injection into mouse zygotes. 200 C57BL/6N zygotes were injected to produce at least one genetically modified mouse. We chose to make this genetic modification on the C57BL/6 background to minimize any strain-dependent effects when we breed these mice with AD mouse models, including the 5xFAD and PS19 lines.

#### Mice

Homozygous ApoE CC mice were cross bred with hemizygous 5xFAD mice to generate hemizygous 5xFAD mice expressing the APOE WT (5xFAD;WT) or heterozygous (5xFAD;Het) or homozygous (5xFAD;Hom) APOE CC variant. Similarly, homozygous APOE CC mice were crossed with hemizygous PS19 mice to obtain hemizygous PS19 mice with WT ApoE (PS19;WT) or heterozygous (PS19;Het) or homozygous (PS19;Hom) APOE CC expression. Finally, homozygous APOE CC mice were crossed with our A53T SynGFP mice (A53T)^35^ to obtain heterozygous (A53T;Het) and homozygous (A53T;Hom) APOE CC mice expressing the A53T α-syn mutation and their respective control mice (A53T;WT). Animals were identified by ear punch right before the weaning and the ear punch was submitted to Transnetyx for genotyping by real-time PCR. For APOE CC genotyping, the introduction of the knock-in mutation removes a restriction enzyme (BstUI) site, making the PCR product resistant to being cut. For all studies, both male and female mice were used.

#### Human

Human tissue was obtained from the Oregon Brain Bank from patients previously seen in the Oregon Alzheimer’s Disease Center (OADC), whose established clinical diagnosis was AD Braak stage VI. Frontal cortex tissue from autopsies of 2 patients was procured in a de-identified manner from the OADC neuropathology core for PHF-tau purification. Tissue use was approved by the IRB at OHSU.

### Method detail

#### Whole genome sequencing and off-target analysis

We performed whole genome sequencing with Genewiz’s, comparing the DNA of a WT and an APOE CC heterozygous mouse to confirm the presence of the mutation in the target site, and without any off-target mutation in encoded regions. Specifically, we performed Illumina HiSeq sequencing technology with 30x coverage of the mouse genome, finding the targeted mutation and no off-target mutations in coding regions.

#### Behavioral analysis

Animals were housed individually beginning a week before the first behavioral test due to the aggressivity of some mice, especially from the 5xFAD mice cohort. The sequence of the behavioral tests was elevated zero maze (day 1), open field (days 2–3), object recognition (days 4-5), and contextual and cued fear conditioning (days 8-9). All procedures were according to the standards of the National Institutes of Health Guide for the Care and Use of Laboratory Animals, followed ARRIVE guidelines, and were approved by the Animal Care and Use Committee of the OHSU. This study contained three studies with the following experimental groups of mice on a C57BL/6J background: 1) 6 months-old 5xFAD/APOE CC Heterozygous (n = 16 mice, [males: n = 8, females: n = 8]), 5xFAD/APOE CC Homozygous (n = 16 mice [males: n = 8, females: n = 8]), and 5xFAD/APOE WT mice (n = 16 mice [males: n = 8, females: n = 8]); 2) 8 months-old PS19/APOE WT (n = 16 mice [males: n = 8, females: n = 8]), PS19/APOE CC Heterozygous (n = 16 mice [males: n = 8, females: n = 8]), and PS19/APOE CC Homozygous (n = 16 mice [males: n = 9, females: n = 7]); and 3) 7 months-old 5xFAD/APOE WT mice injected with tau (n = 7 mice [males: n = 4, females: n = 3]); 5xFAD/APOE CC Homozygous mice injected with tau (n = 7 mice [males: n = 4, females: n = 3]).

#### Nest building

Nest building was assessed Wednesday-through Friday prior to behavioral tests using an established protocol.^49^ Briefly, mice were housed in a clean cage and provided 2 pressed cotton squares. Photos of the home cage were taken 48 h later and visually rated by a blinded scorer on a 5-point nest rating scale.

#### Elevated zero maze

Mice were placed in a custom-built elevated zero maze consisting of two enclosed areas and two open areas (6 cm wide). Mice were placed in the closed part of the maze and allowed free access for 10 min. A video tracking system (set at six samples per second) was used to calculate the distance moved, velocity, and time spent in the open and closed areas. Mice that are anxious in the elevated zero maze spend less time in the open areas.^50^

#### Open field and novel object recognition

The mice were put in an open field enclosure (16 x 16 inches) for 10 min on two subsequent days. On day 3, the open field contained two identical objects for a 15-minute trial. The next day, one object was replaced with a novel object for a 15-minute trial. Between trials, the arenas and objects were cleaned with 0.5% acetic. Interaction within a 2 cm proximity with the object was coded as object exploration by hand scoring videos acquired with Noldus Ethovision software. The percent time spent with the novel object (time spent exploring the novel object/(time spent exploring the familiar and novel objects) in the first 5-min bin of the test was analyzed to assess object recognition. The outcome measures in the open field analyzed were: 1) distance moved in the open field in the absence and presence of objects, an activity measure; 2) the difference in the distance moved in the open field over days, habituation to the open field, a cognitive measure; 3) time spent in the center of the open field, an anxiety measure; and 4) the percent time spent exploring the novel object on the second day of open field testing in the presence of objects, a cognitive measure.

#### Contextual and cued fear conditioning

On the first day of the conditioned fear test, each mouse was placed in a fear conditioning chamber and allowed to explore it for 1.5 minutes before the delivery of a 30 second tone (80 dB) which was immediately followed by a 2 second foot shock (0.35 mA). Ninety seconds later, a second tone-shock pair was delivered. Mice were removed from the testing chambers 90 seconds after the second shock and were returned to their home cages. Chambers were cleaned with 0.5% acetic acid between animals. The pre-tone time, which was the first 2 minutes of the trial, was used as the baseline measure for freezing behavior. Mice were removed from the testing chambers 10 seconds after the end of the second shock and returned to their home cages. On day 2, each mouse was first placed in the fear conditioning chamber containing the exact same context but without delivery of a tone or foot shock. Freezing was analyzed for 5 minutes. The context of the chambers was changed by adding a smooth floor texture over the grid floor, inserting the shape of a triangle, adding a new scent (hidden vanilla soaked nestlets), and by cleaning the chamber with 70% ethanol rather than acetic acid. Two-three hours after the last contextual test for each mouse, the mice were assessed for cued fear conditioning. Mice were placed in the chambers containing the modified context and were allowed to explore for 90 seconds before they were re-exposed to the fear conditioning tone for 3 minutes. Freezing behavior was analyzed for the first and last 3 minutes of the cued fear conditioning test. Freezing was measured using a motion index, calculated based on a proprietary motion analysis algorithm in the Med Associates Video Freeze Software. Briefly, the software analyzes and acquires videos of the trials at a frequency of 30 frames/second. The motion index is based on the sum of the pixel changes in a frame compared to those of a reference frame and to those of successive frames. The reference frame is based on a video capture when the mouse is not in the chamber. The motion index threshold used in the current study was 18. This means that the motion index had to remain below 18 pixels changes to be considered freezing. Analysis of group differences in freezing before delivery of the first tone during fear conditioning training on day 1 were analyzed as measure of baseline freezing. This allowed us to determine whether there were potential pre-conditioning group differences in behaviors such as immobility, which could contribute to freezing scores. Potential group differences in the motion index during the two shocks on day 1 were also analyzed. This allowed us to determine whether there were possible group differences in sensory response to the shocks. Finally, the freezing percentage during contextual and cued testing on day 2 were analyzed.

#### Plasma extraction and lipid analysis by fast protein liquid chromatography

Whole blood (∼75-100 mL) was collected from the retro-orbital sinus from mice under isoflurane anesthesia using heparin coated microhematocrit capillary tubes (Fisher Scientific, Hampton, NH). Blood was immediately transferred to microcentrifuge tubes containing 0.5 M ethylenediaminetetraacetic acid (EDTA) and gently mixed to prevent clotting. Microcentrifuge tubes containing EDTA-treated whole blood were centrifuged at 12,000 rpm for 5 minutes and the plasma supernatant was collected for lipid assessments. Plasma was pooled from 3 mice/sex/age group and subjected to fast protein liquid chromatography (FPLC) to fractionate the plasma lipoproteins based on size. The total cholesterol (mg/dL) in each FPLC fraction was determined using cholesterol liquid reagents by measuring absorbance at 490 nm with a SpectraMax iD3 microplate reader. The total triglyceride (mg/dL) in each FPLC fraction was determined using triglycerides liquid reagents by measuring absorbance at 540 nm with a SpectraMax iD3 microplate reader.

#### PHF-tau purification from postmortem human AD brains and stereotactic injections

Following the protocol developed by *Guo et al*.,^35^ 6 grams of gray matter from the frontal cortex of two de-identified human AD patient brains (Braak stage VI) with a postmortem interval inferior to 24h (Oregon Brain Bank, USA) were homogenized using a Dounce homogenizer in nine volumes (v/w) of high-salt buffer (10 mM Tris-HCl, pH 7.4, 0.8M NaCl, 1mM EDTA, and 2mMdithiothreitol (DTT), with protease inhibitor cocktail, phosphatase inhibitor, and PMSF) with 0.1% sarkosyl and 10% sucrose added and centrifuged at 10,000 g for 10 min at 4°C. Pellets were reextracted twice using the same buffer, and the supernatants were filtered and pooled. Later, sarkosyl was added to reach 1% and samples were rotated for 1h at room temperature for proper mixing. Then, samples were centrifuged at 300,000 g for 60 min at 4°C. Pellets containing pathological tau were washed once in PBS and resuspended in 600 µL PBS by passing through 27-G 0.5-in needle. Samples were briefly sonicated (20 pulses at 0.5 s/pulse) and centrifuged again at 100,000 g for 30 min at 4°C. Pellets were resuspended in 200 µL PBS, sonicated again using the same settings as before, boiled at 95°C for 10 min and centrifuged at 10,000 g for 30 min at 4°C. Final supernatants, containing AD paired helical filaments (PHF) of tau and referred to as PHF-tau, were frozen at −80°C until they were used for intracranial injections. Samples were diluted to a final concentration of 0.4 µg/µL with PBS and sonicated right before the injection. 4-month-old 5xFAD;WT or 5xFAD;Hom mice were deeply anesthetized with 1-2% isoflurane and immobilized in a stereotactic frame. 2 µg of PHF-tau (1 µg/site) were unilaterally injected first into the dorsal hippocampus (bregma: −2.5 mm; lateral: 2 mm; depth: −2.4 mm from the skull) and then into the overlaying cortex (bregma: −2.5 mm; lateral: 2 mm; depth: −1.4 mm from the skull) using a Hamilton syringe. Mice were monitored for any discomfort for the next 3 days after the surgery.

#### Intracortical and hindlimb gastrocnemius muscle α-syn PFF injections

Mouse α-syn PFF injections were done according to published protocols for PFF generation and preparation.^37,51,52^ For intracortical injections, 4-month-old Heterozygous and homozygous APOE CC mice and their respective control mice were anesthetized with 1-2% isoflurane and immobilized in a stereotactic frame. 5 µg (2.5 µL) of freshly sonicated mouse α-syn PFF was injected into right hemisphere primary sensory cortex (bregma: −1.5 mm; lateral: 1 mm; depth: −0.3 mm from the skull). For intramuscular injections, 10 µg (5 µL) of freshly sonicated mouse α-syn PFF was injected into the gastrocnemius muscle of 1-2%-anesthetized A53T mice heterozygous (A53T;Het) or homozygous (A53T;Hom) APOE CC mice and their respective control mice (A53T;WT).

#### Tissue preparation for histological analysis and immunohistochemistry

When the desired age was reached, mice were euthanized via CO2 asphyxiation and decapitation. Whole brain and spinal cord were dissected, placed in scintillation vials with 10% formalin for 48h and then stored in PBS containing sodium azide (0.05%) until use. After dehydration, tissue was embedded in paraffin and serial 5-µm-thick coronal sections were cut through the entire brain using a conventional microtome. Sequential sections (1 mm apart) were immunostained covering the anterior-posterior axis. In brief, brain or spinal cord slices were deparaffinized for at least 2h at 70°C in the oven, followed by xylene immersion and rehydration in graded ethanol solutions to distilled water. After, antigen retrieval was performed by first, 5 min immersion in acetic acid at room temperature and a 45 min immersion in citrate buffer (10 mM Sodium citrate, 0.05% Tween 20, pH 6.0) at 85°C. Slides were then blocked in 3% milk-PBS solution followed by an 4°C overnight incubation with the following primary antibodies: Aβ (4g8, 1:5000), phosphorylated tau (AT8, 1:1000) aggregated p-α-syn (pS129, 1:5000 in non-transgenic tissue, 1:40000 in A53T tissue). The next day, slides were washed with PBS and incubated with a horseradish peroxidase-conjugated secondary antibody for 1h, followed by a 1h incubation in ABC solution. Finally, sections were developed and stained using the peroxidase substrate 3,3’-Diaminobenzidine (DAB) for 10 min, washed in water for 5 min, counterstained with hematoxylin and dehydrated with ethanol and xylene before mounting the coverslip using Cytoseal 60 as the mounting media.

#### Tissue preparation for biochemical analysis and Enzyme Linked-Immuno-Sorbent Assay (ELISA)

Mice were euthanized via CO2 asphyxiation and decapitation. Brains were extracted and the cortex and the hippocampus of each brain hemisphere were micro-dissected and flash frozen. Tissue was homogenized in eight volumes (v/w) of PBS with protease and phosphatase inhibitor cocktails and centrifuge at 5,000 g for 5 min at 4°C. Supernatant was ultracentrifuge at 125,000 g for 1h at 4°C. The supernatants were saved as the PBS-soluble (soluble) samples, while the pellets were resuspended and homogenized in 5M guanidine-HCL using the same volume as before and briefly sonicated (20 pulses at 0.5 s/pulse). Then, the homogenates were mixed on an orbital shaker at room temperature for at least 3h and ultracentrifuged at 200,000 g for 1h at 4°C. Finally, supernatants were saved as the guanidine-soluble (insoluble) fraction, containing aggregated proteins. Quantification of Aβ40 and Aβ42 protein levels in the soluble and insoluble fractions in the cortex and the hippocampus of 5xFAD;WT, 5xFAD;Het and 5xFAD;Hom mice were acquired using the Human Aβ40 and Human Aβ42 ELISA kits according to the manufacturer’s guidelines. Quantification of pS396 tau and total tau protein levels in the soluble and insoluble fractions in the cortex and hippocampus of PS19;WT, PS19;Het and PS19;Hom mice were acquired using the Human Tau [pS396] and Human Tau Total ELISA kits following the manufacturer’s instructions.

#### Image acquisition

For analysis purposes, brightfield images of slides containing brain slices were obtained using Zeiss Axioscan 7 using a 10x 0.45NA Plan-Apochromat objective. This system autofocuses by searching for a pre-defined z-stack and determines the plane with the highest contrast in brightfield. Representative images for figures were obtained using an Axio Imager Z2 with an Apotome™ attachment and two cameras: an AxioCam 512c and an AxioCam 506 mono. Images were taken using a Plan-Apochromat 10x/0.45 WD=2.0 or 20x/0.8 WD=0.55 objectives. Both systems are driven by ZEN (v3.12 and v3.11; Carl Zeiss) and are part of the Advanced Light Microscopy Core at the Jungers Center, OHSU, Portland, Oregon.

### Quantification and statistical analysis

#### FIJI ImageJ and Visiopharm analysis

All the stainings, except for the AT8 staining in PHF-tau injected 5xFAD mice, were analyzed using Fiji ImageJ (NIH). In brief, images were deconvoluted by using the Color Deconvolution plugin to split the DAB signal from the hematoxylin and the background. DAB images were then converted to 8-bit gray-scale and adjusted for a threshold to further distinguish signal from background. Total number of Aβ plaques, tau aggregates or α-syn inclusions were quantified using the analyze particle feature. Image analysis of the AT8 staining in the PHF-tau injected 5xFAD was performed using Visiopharm software. A deep learning classifier was trained to first detect the tissue and then to detect the AT8 positive aggregates. In brief, a tissue region of interest was created after training two variables (tissue and non-tissue regions). Another deep learner classifier was developed and trained for detecting tau positive signal after training two variables (tau and background). Abnormal signals, such as tissue folds or bubbles, were excluded from the analysis.

#### Statistical analysis

All data are reported as mean ± standard error of the mean and were analyzed using SPSS or GraphPad software. Genotype and sex or genotype and age were included as factors in analysis of variance (ANOVAs). Mantel-Cox and Gehan-Breslow-Wilcoxon tests were used for survival analysis. Unpaired t-test was used when only two groups were compared. Pearson correlation coefficient was calculated for correlation analysis between pathological levels or between pathological levels and behavior. Repeated measures ANOVAs were used when appropriate during behavioral tests. When the standard deviations (SD) were different among the groups, Brown-Forsythe ANOVA and Welch’s ANOVA tests were used. Statistical significance was considered as *p* < 0.05. The significant level is indicated using asterisks: * *p* < 0.05, ** *p* < 0.01, *** *p* < 0.001, **** *p* < 0.0001. When sphericity was violated (Mauchly’s test), Greenhouse-Geisser corrections were used. For analysis of object recognition, paired t-tests were used for each experimental group. All researchers were blinded to genotype and sex, and the code was only broken after the data were analyzed.

## Supplemental information

**Supplementary Table 1. Comparison between previous reports and this work exploring the effects of APOE CC.**

**Supplementary Figure 1. Behavioral analysis of 6 months-old 5xFAD mice with heterozygous or homozygous murine APOE CC expression and their respective control mice.** Open field (Day 1 speed: sex effect: F (1, 42) = 10.80, *p* = 0.0021; Day 2 distance moved: sex: F (1, 42) = 16.42, *p* = 0.0002; and genotype effects: F (2, 42) = 6.049, *p* = 0.0049; speed: sex: F (1, 42) = 16.29, *p* = 0.0002; and genotype effects: F (2, 42) = 6.161, *p* = 0.0045; time in center: sex effect: F (1, 42) = 5.283, p = 0.0266), novel object recognition (NOR. Day 1 speed: sex F (1, 42) = 22.11, *p* < 0.0001; and genotype effects: F (2, 42) = 6.971, *p* = 0.0024; Day 2 distance moved: sex: F (1, 42) = 17.60, *p* = 0.0001; genotype: F (2, 42) = 7.131, *p* = 0.0022; and interaction effects: F (2, 42) = 3.285, *p* = 0.0473; speed: sex: F (1, 42) = 17.26, *p* = 0.0002; genotype: F (2, 42) = 7.042, *p* = 0.0023; and interaction effects: F (2, 42) = 3.249, *p* = 0.0487), elevated zero maze (EZM) and cued (Baseline freezing: sex effect: F (1, 42) = 4.730, *p* = 0.0353), and contextual (Freezing: sex effect: F (1, 42) = 5.737, *p* = 0.0211; Distance moved: sex effect: F (1, 42) = 9.450, *p* = 0.0037) fear conditioning (FC. Train: Tone 2 freezing: sex effect: F (1, 42) = 5.594, *p* = 0.0227; Tone 2 distance moved: sex effect: F (1, 42) = 6.967, *p* = 0.0116; ISI2 distance moved: genotype effect: F (2, 42) = 4.799, *p* = 0.0133; Shock 1 distance moved: interaction effect: F (2, 42) = 3.395, *p* = 0.0430) were performed. Two-way ANOVA followed by Tukey’s post hoc was used as a statistical test. Data represents the speed (cm/s), distance moved (cm), number of times, time in and latency to the center (s) and percentage of time/freezing ± SD. * *p* < 0.05, ** *p* < 0.01, *** *p* < 0.001, **** *p* < 0.0001.

**Supplementary Figure 2. Behavioral analysis of 8 months-old PS19 mice with heterozygous or homozygous murine APOE CC expression and their respective control mice.** Nest building, open field (Day 1 distance moved: sex effect: F (1, 42) = 11.40, *p* = 0.0016; speed: sex effect: F (1, 42) = 12.17, *p* = 0.0012; latency to center: genotype effect: F (2, 42) = 4.174, *p* = 0.0222; Day 2 latency to center: genotype effect: F (2, 42) = 5.493, *p* = 0.0076), novel object recognition (NOR. Latency to object 2: interaction effect: F (2, 42) = 4.392, *p* = 0.0185; time in objects: genotype effect: F (2, 42) = 3.259, *p* = 0.0483), elevated zero maze (EZM) and cued and contextual fear conditioning (FC. Train: Tone 1 distance moved: sex effect: F (1, 42) = 4.980, *p* = 0.0310; Tone 2 freezing: sex effect: F (1, 42) = 4.169, *p* = 0.0475; ISI1 distance moved: sex effect: F (1, 42) = 6.529, *p* = 0.0143) were performed. Two-way ANOVA followed by Tukey’s post hoc was used as a statistical test. Data represents the score, speed (cm/s), distance moved (cm), number of times, time in and latency to the center (s) and percentage of time/freezing ± SD. * *p* < 0.05, ** *p* < 0.01.

**Supplementary Figure 3. Behavioral analysis of 7 months-old PHF-tau injected 5xFAD;WT and 5xFAD;Hom mice.** Open field (Day 1 distance moved: genotype effect: F (1, 10) = 8.845, *p* = 0.0139; speed: genotype effect: F (1, 10) = 8.964, *p* = 0.0135; times in center: genotype effect: F (1, 10) = 7.811, *p* = 0.0190), novel object recognition (NOR), elevated zero maze (EZM. Time and %time in open arms: interaction effect: F (1, 10) = 10.66, *p* = 0.0085) and cued and contextual fear conditioning (FC. Train: Baseline distance moved: sex: F (1, 10) = 10.54, *p* = 0.0088; and interaction effects: F (1, 10) = 4.973, *p* = 0.0498) were performed. Two-way ANOVA followed by Tukey’s post hoc was used as a statistical test. Data represents the speed (cm/s), distance moved (cm), number of times, time in and latency to the center (s) and percentage of time/freezing ± SD. * *p* < 0.05, ** *p* < 0.01.

