## Supplementary material for "The Christchurch point mutation in mouse APOE reduces Aβ-induced tau and α-synuclein pathologies": Supplemetary Table and Figures

### Supplementary Table 1

| Study | System | Plasma lipids | Aβ | Tau | Aβ-induced tau | α-syn | Behavior | Sex differences |
|---|---|---|---|---|---|---|---|---|
| Nelson et al., 2023 | Human APOE4 CC | × | × | Reduction in PS19 | × | × | × | × |
| Chen et al., 2024 | Human APOE3 CC | Increased cholesterol | Reduction in APP/PSEN1 | No effects after PHF-tau injection | Reduction after PHF-tau injection in APP/PSEN1 | × | - Y maze<br>- Cued fear conditioning | × |
| Naguib et al., 2025 | Human APOE3 CC | × | × | Reduction in PS19 | × | × | × | × |
| Tran et al., 2025 | Murine APOE CC | Increased cholesterol | Reduction in 5xFAD | No effects in PS19 | × | × | - Hindlimb clasping<br>- Elevated plus maze<br>- Open field | × |
| Gunaydin et al., 2024 | AAV human APOE2 CC | × | Reduction in APP/PSEN1/TRE4 | Reduction in PS19/TRE4 | × | × | - Nest building<br>- Y maze<br>- Novel object recognition<br>- Barnes maze | No |
| Chen et al., 2025 | AAV human APOE3 CC | × | Reduction in 5xFAD | Reduction in PS19 | Reduction after recombinant tau PFF injection in 5xFAD | × | - Morris water maze<br>- Contextual and cued fear conditioning | × |
| Soto-Faguás et al., 2025 (This report) | Murine APOE CC | Increased cholesterol and triglyceride | Reduction in 5xFAD females | No effects in PS19 | Reduction after PHF-tau injection in 5xFAD | Reduction in intracortical and intramuscular α-syn PFF injection | - Nest building<br>- Elevated zero maze<br>- Open field<br>- Novel object recognition<br>- Contextual and cued fear conditioning | Yes |

Supplementary Fig. 1

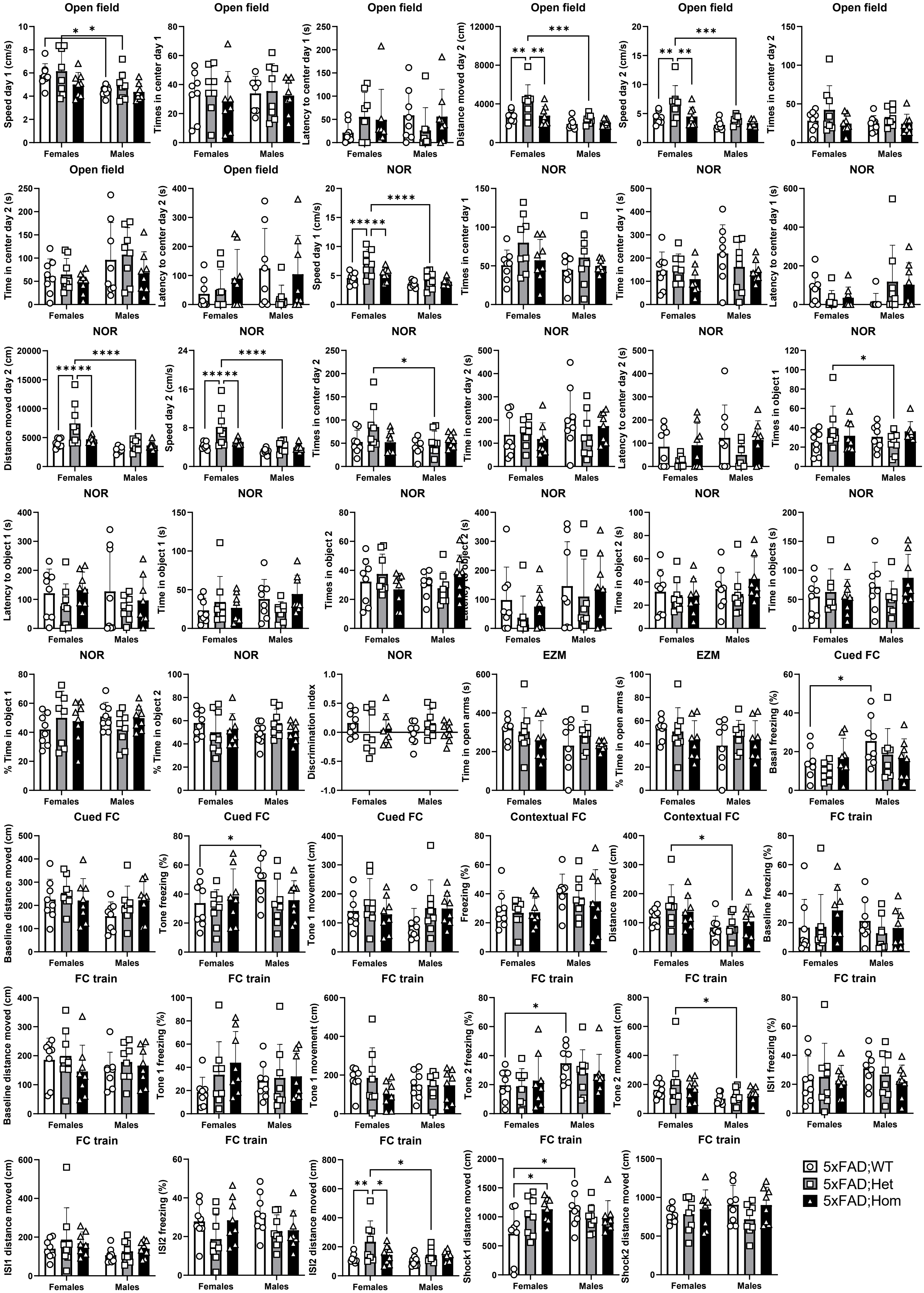

Supplementary Fig. 2

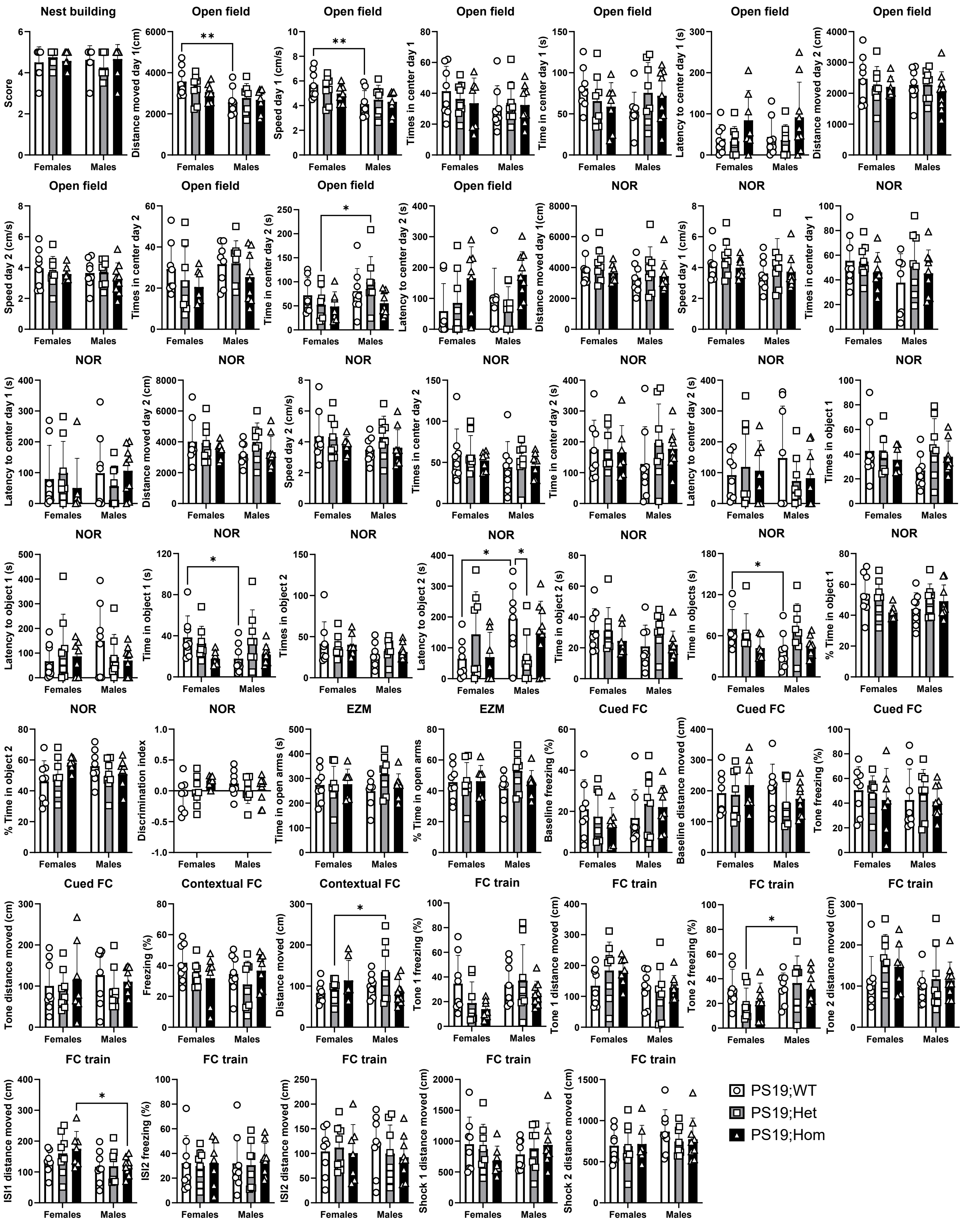

### Supplementary Fig. 3

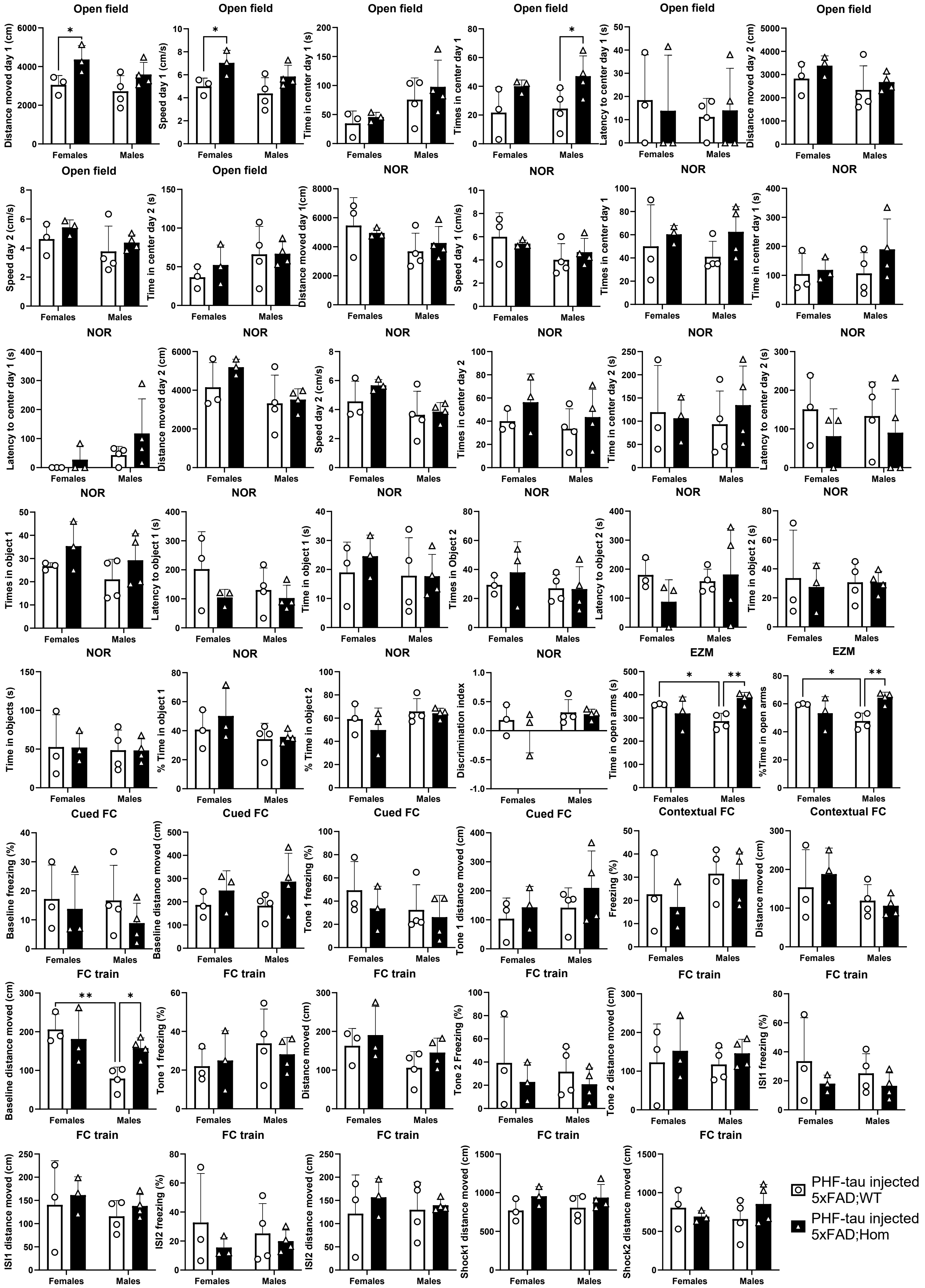
